# PolyFold: Homology Filtering Approach Alters Apparent Structure Prediction Model Performances

**DOI:** 10.64898/2026.06.16.732304

**Authors:** Henry Stephenson, Dan Voicu, Victor Novakov, Michelle Levy, Joseph Marsilio, Christopher Kottke, Sina Sarparast, Maximilian Ebert

## Abstract

As structure prediction models become increasingly central to biological research and drug discovery, rigorous and reproducible benchmarking is critical to assess model performance and generalization. PolyFold is an end-to-end workflow automation and benchmarking framework that streamlines input file creation, batch automation, and a more conservative evaluation of model performances. We conduct a head-to-head comparison of Boltz-2 and OpenFold 3 – leading open-source alternatives to AlphaFold 3. We first curate an evaluation set of several thousand high-quality PDB structures, homology-filtered against both models’ training sets, then automate inference and evaluation using PolyFold. Using a cluster analysis, we directly connect features of the underlying structures to model performance profiles, demonstrating the nonrandom nature of failure and success modes. While initial analyses using conventional homology filtering techniques show performance differences favouring Boltz-2, we show that using minimum chain fractional identity for novelty filtering can cause residual training-data leakage in multichain complexes, inflating apparent performances. To address this, we introduce a length-weighted average fractional identity (WAFI) for whole-complex homology filtering. This provides a more conservative evaluation by reducing residual leakage. While Boltz-2 maintains higher aggregate metric averages for categories such as nucleic acid-containing structures, membrane-bound proteins, and antibody–antigen complexes, the performance gap significantly narrows or disappears for effectively all partitions after implementing WAFI filtering, demonstrating the need for more stringent conventions in homology filtering. Both models still struggle with homomultimers, small molecule realism, and structures with non-canonical amino acids, demonstrating persisting limitations in the field. In summary, PolyFold enables practitioners to rapidly compare models on a variety of inference tasks and supports the broader adoption of accessible computational tools to facilitate biomolecular science.

## Introduction

The biomolecular folding problem has been a foundational challenge in the biological and life sciences for more than 50 years (Jumper et al., 2021; Malhotra et al., 2025; Dill et al., 2008). Consistent and accurate computational prediction of three-dimensional biomolecular structures based solely on the primary sequence would overcome the massive bottleneck of experimental structure determination, a laborious, expensive process requiring access to advanced laboratory equipment and specially trained personnel (Jumper et al., 2021; Qiu et al., 2024). As such, biomolecular structure prediction models have the potential for significant impact on the rate of innovation by expediting development of life-saving medicines, addressing critical environmental concerns, and combatting infectious diseases (Qiu et al., 2024; Kovalevskiy et al., 2024; Nji et al., 2025; Fang et al., 2025; Gong et al., 2025).

Since the release of the first AlphaFold model in 2018, Google Deepmind has played a pivotal role in *in silico* structure prediction, achieving the highest overall performance on the 13th and 14th CASP experiments in 2018 and 2020 respectively (Deepmind has not entered the competition more recently) (Jumper et al., 2021; Malhotra et al., 2025; Abbass, 2025). While AlphaFold 3 shows promise in advancing structural biology, drug discovery, and protein engineering, AlphaFold 3’s prohibited use policies present barriers to widespread application, especially in commercial settings, limiting its utility for biotechnological and pharmaceutical advancement (Jumper et al., 2021; Malhotra et al., 2025; Abbass, 2025). This has led to the development of a suite of alternatives to AlphaFold, including Boltz (Wohlwend et al., 2025; Passaro et al., 2025), Chai (Boitreaud et al., 2024, 2025), ESMFold (Lin et al., 2023), Open-Fold (Ahdritz et al., 2024; The OpenFold3 Team, 2025), Protenix (Chen et al., 2025; Zhang et al., 2026), and RoseTTAFold (Corley et al., 2025), among others (Wu et al., 2022; Liu et al., 2024; Mirdita et al., 2022). Despite competitive performance for state-of-the-art openuse algorithms, AlphaFold 3 still outclasses even the latest models in nearly all folding tasks (Passaro et al., 2025; The OpenFold3 Team, 2025). Some models approach AlphaFold’s results on monomer folding or protein-RNA interface preservation, though for other key interactions – such as antibody-antigen complexes – the differential is pronounced (see Figure 1).

**Figure 1.**
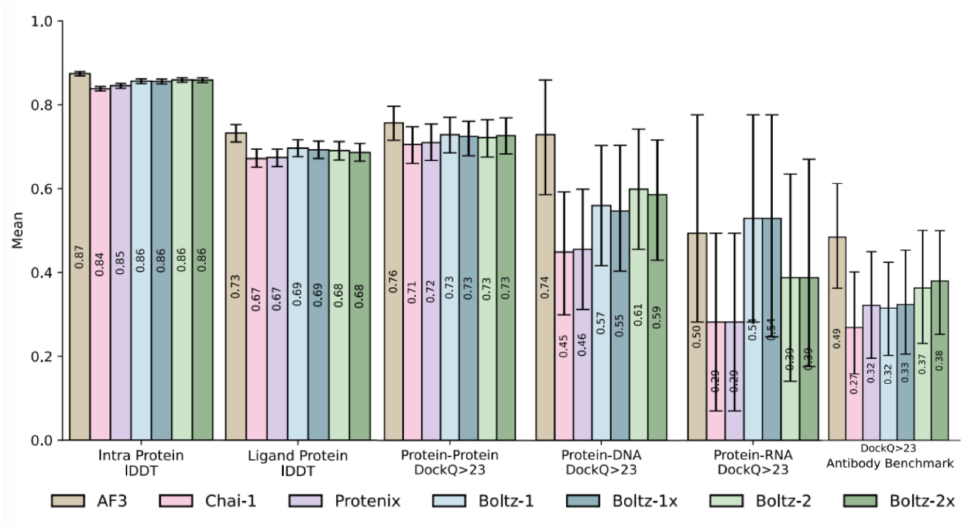
Comparing AlphaFold 3, Boltz models, and others’ performances on key folding tasks. From left to right: mean lDDT within a single protein chain, ligand-protein interface lDDT, proportion of protein-protein interfaces with DockQ > 0.23 (DockQ > 0.23 is classed as “acceptable” interface preservation), protein-DNA interfaces with DockQ > 0.23, protein-RNA interfaces with DockQ > 0.23, and proportion of antibody-antigen interfaces with DockQ > 0.23 on an undisclosed challenging antibody benchmark discussed in Passaro et al., 2025 (Figure modified from Passaro et al., 2025).

In addition to underperformance of open-use models, major questions persist on the utility of protein structure prediction algorithms in specific folding tasks, such as intrinsically disordered proteins (Haynes et al., 2006; Liu et al., 2006; Dyson, 2023), conformationally dynamic proteins (amyloidogenic proteins, GPCRs, ion channels, etc.) (Tuinstra et al., 2008; Hoffmann et al., 2008; Bhattacharya et al., 2008; Zhang et al., 2024; Sanganna Gari et al., 2021; Nekouzadeh and Rudy, 2016; Li and Liu, 2022; Xu et al., 2005; Pinzi et al., 2023; Lee et al., 2025), highly variable interfaces (such as antibody complementary-determining regions (CDRs), TCR-pMHC interactions, etc.) (Blevins et al., 2016; Spoendlin et al., 2025), post-translationally modified proteins (e.g. lectins, glycoproteins, phosphorylated or acetylated proteins, etc.) (Huang and Liu, 2024; Medvedev et al., 2025), and protein complexes with non-protein components (RNA-protein complexes, DNA-transcription factor interactions, small-moleculebinding proteins, metalloenzymes, etc.) (Varadi et al., 2023; Abramson et al., 2024; Krokidis et al., 2025). Figure 1 demonstrates a stark difference in performance for single-chain protein structure prediction (which produces high-performing metrics for AlphaFold 3 as well as most leading open-use models) compared to tasks such as protein-DNA, protein-RNA, and protein-smallmolecule complexes. While general protein-protein interfaces have relatively high success in terms of interface preservation, specific cases like antibody-antigen complexes have reduced performance in terms of interface lDDTs and correct prediction of specific sidechain interactions (Passaro et al., 2025). These results raise uncertainty about the ability for structure prediction models to provide sufficiently accurate outputs in these essential use cases, bringing attention to the current limitations that must be overcome to make ML-based biopolymer folding a consistently useful and trustworthy tool for scientifically rigorous applications.

In this work, we develop a comprehensive evaluation pipeline, PolyFold, which automates file preparation, model inference, and analysis of predicted structures through an easy-to-use workflow automation tool (WAT), streamlining head-to-head evaluations of structure prediction algorithms. Using PolyFold, we perform an in-depth comparison of two leading latest-generation open-use models: Boltz-2 and OpenFold 3. Crucially, our evaluation reveals that varying the strategy or stringency of homology filtering during dataset curation significantly affects apparent performance differentials between the models, or can even change whether a significant performance gap exists, highlighting limitations in conventional filtering practices when attempting to manage data leakage.

PolyFold is complementary to existing benchmarking infrastructures and past comparative studies. While frameworks like FoldBench evaluate a broad panel of all-atom predictors across multiple biomolecular tasks (Xu et al., 2025), and the OpenStructure suite provides comprehensive automated structural comparison and scoring (Studer et al., 2026), PolyFold integrates dataset curation, model-specific inference, and downstream comparative analysis. Furthermore, whereas initial technical reports offered brief, aggregate comparisons of Boltz-2 and OpenFold 3 across limited partitions (The OpenFold3 Team, 2025), our study uses PolyFold to conduct a focused, task-specific evaluation that explicitly demonstrates how complex-level homology filtering choices alter apparent model performance.

By evaluating performance across specific folding tasks and interaction classes — rather than relying on aggregate measures that can mask task-specific weaknesses — PolyFold enables researchers to determine the model most appropriate for their specific biological problem, saving computational resources and setup time. In addition to providing an independent, thirdparty perspective on the relative capabilities of Boltz-2 and OpenFold 3 under multiple filtering stringencies, we provide open access to a curated dataset with transparent filtering logic to serve as a clean resource for future model development. Future work may leverage Poly-Fold to expand these head-to-head evaluations to additional models, such as AlphaFold, Chai, ESMFold, etc. and incorporate increasingly specialized data partitions.

## Results and Discussion

### Conventional Homology Filtering Suggests a Significant Performance Differential

To evaluate OpenFold 3 and Boltz-2 performances, we compute several metrics. Mathematical and qualitative descriptions of all metrics used in this work can be found in the Evaluation Metrics subsection of Methods. We begin with a general comparison of both models using four structural metrics: TM-score, mean lDDT, mean ilDDT, and mean DockQ. Table 1 summarizes these metrics, their definitions, and their score ranges. All four metrics range from 0 to 1, with higher scores indicating better structural agreement or quality.

**Table 1.**
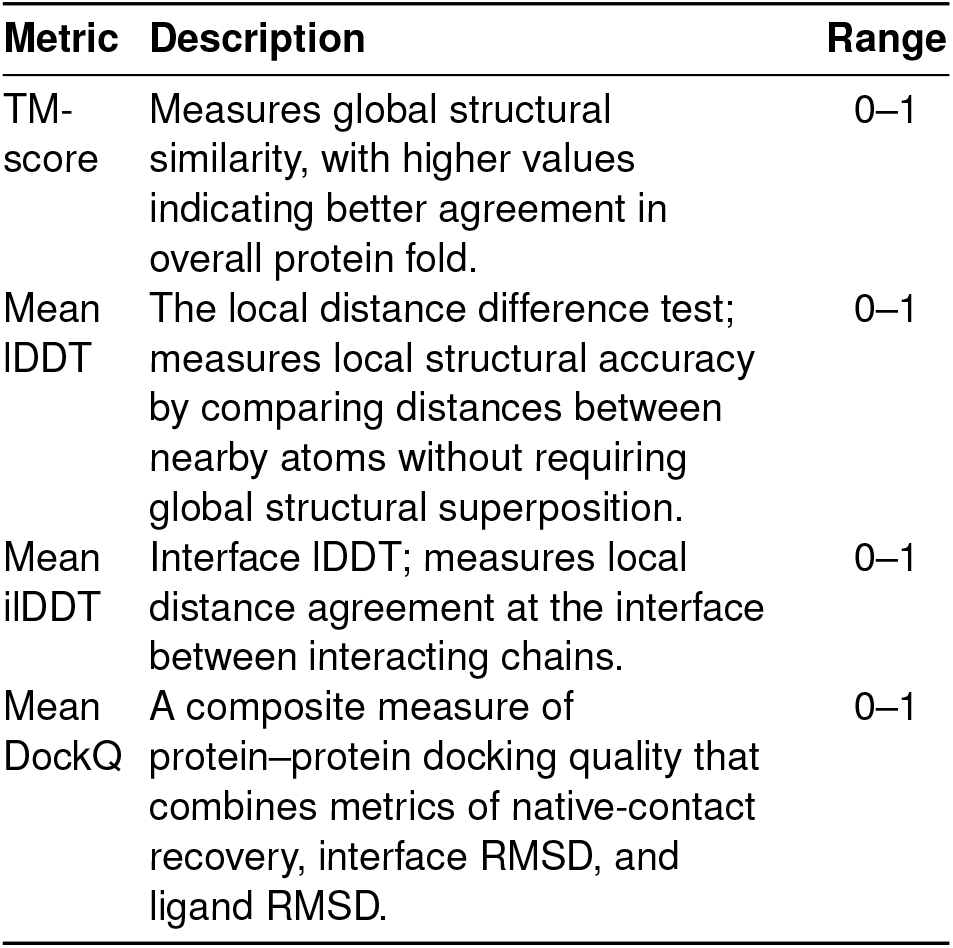
Structural quality metrics for comparing OpenFold 3 and Boltz-2.

| **Metric** | **Description** | **Range** |
|---|---|---|
| TM-score | Measures global structural similarity, with higher values indicating better agreement in overall protein fold. | 0–1 |
| Mean lDDT | The local distance difference test; measures local structural accuracy by comparing distances between nearby atoms without requiring global structural superposition. | 0–1 |
| Mean ilDDT | Interface lDDT; measures local distance agreement at the interface between interacting chains. | 0–1 |
| Mean DockQ | A composite measure of protein–protein docking quality that combines metrics of native-contact recovery, interface RMSD, and ligand RMSD. | 0–1 |

To address information leakage, we first apply a relatively lenient novelty filter, namely a fractional identity (F_ident_) cutoff of 0.6, to all structures in the evaluation set before comparing Boltz-2 and OpenFold 3 performances (see Methods for further discussion on assembly of the evaluation set as well as F_ident_). Using this inclusion approach, Boltz-2 has higher average scores than OpenFold 3 for all metrics, with the percentage of data points favouring Boltz-2 ranging from 59.5% for mean lDDT to 66.9% for mean DockQ (Figure 2). The distribution of data points also demonstrates a larger mean distance from the diagonal for data points where Boltz-2 scores higher than OpenFold 3 than for data points showing the opposite trend.

**Figure 2.**
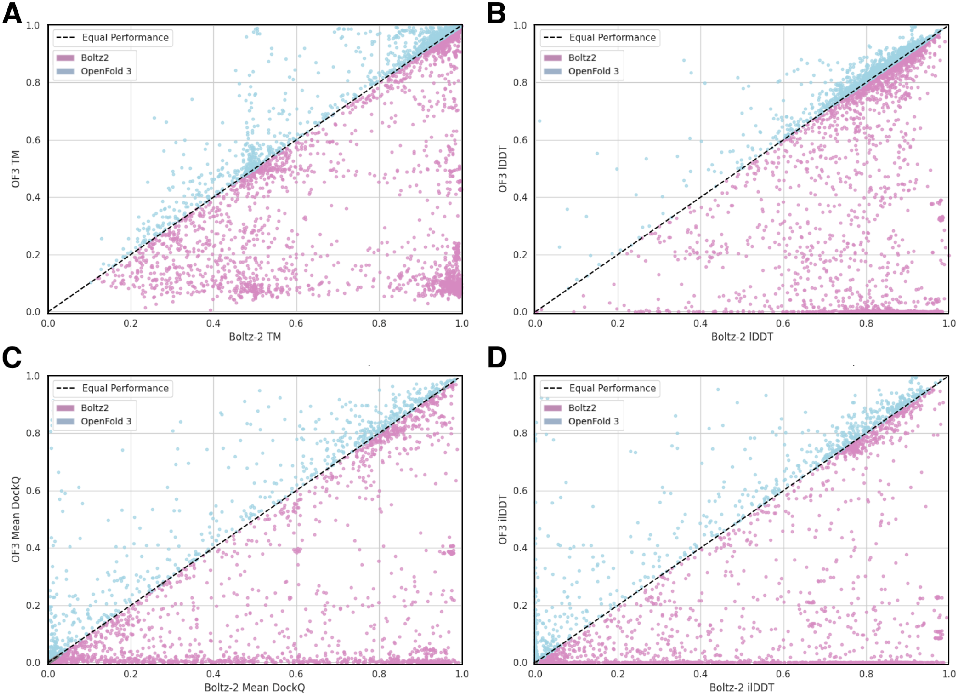
General comparison of Boltz-2 vs. OpenFold 3. (A) TM-scores: 62.5% of samples have higher Boltz-2 scores. (B) Mean lDDTs: 59.5% of samples have higher Boltz-2 scores. (C) Mean DockQs: 66.9% of samples have higher Boltz-2 scores. (D) Mean ilDDTs: 65.8% of samples have higher Boltz-2 scores. The provided rates for each plot are the counts of data points below the dashed diagonal, representing the line of equal performance. Inclusion on the plot requires *≥* 1 chain in a structure having fractional identity, or Fident, less than or equal to 0.6 compared to the union of Boltz-2 and OpenFold 3 training sets. In other words, there must be at least one polymeric chain with a certain degree of novelty compared to training examples. See Methods for further discussion of F_ident_.

Higher general scores for Boltz-2 over OpenFold 3 for interface metrics (e.g., ilDDT and DockQ) may additionally indicate that Boltz-2 has a particular advantage for predicting biomolecular complexes (while the differential performance is more mild for structures with a single polymer chain). However, this analysis merely demonstrates higher average metrics for Boltz-2 under a minimally rigorous, yet conventional, novelty filter; results presented later in this work demonstrate a notable limitation of this approach that may lead to inequitable inflation of model performances. These results thus serve as a baseline to which downstream sections compare using more stringent homology-filtering approaches, rather than suggesting a *bona fide* performance differential of this magnitude.

### Clustering Reveals Relationships Between Structural Features and Model Performances

To interpret the relationship between structural patterns and model performance profiles, we performed densitybased clustering by vectorizing examples based on their performance metrics. That is, each dimension in the vector represents a Z-score-normalized performance metric – e.g., RMSDs, lDDTs, DockQs, etc. – of either the Boltz-2 or OpenFold 3 output structure compared to ground truth (See the HDBSCAN clustering subsection in Methods). High-density clusters were discovered in this high-dimensional space, indicating consistent prediction patterns based on input data sharing similar features (see Figure 3 for cluster divisions along key 2D axes).

**Figure 3.**
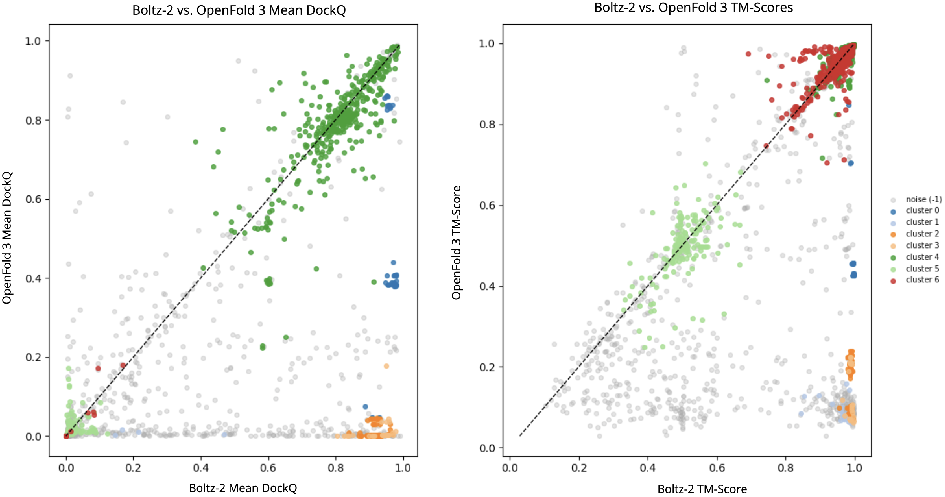
Clustering along key metric axes. Comparison of Boltz-2 and OpenFold 3 along mean DockQ (left) and TM (right) with HDBSCAN clusters annotated shows distinct performance regimes across the clusters.

For each cluster, we identified which features of the underlying structure (e.g., sequence length, presence of small molecules, number of chains, minimum fractional identity (F_ident_) to training set examples, etc.) were associated with the cluster; we performed effect size analysis on each cluster, comparing in-cluster character to out-of-cluster (i.e., the remaining examples) character to identify the most significant features distinguishing particular clusters (see Appendix D in the Supplementary Information). We additionally performed normalized multivariate regressions to construct models for the features driving specific performance profiles for a given cluster (Appendix D). Altogether, we identified several clusters characterized by features such as: single chains in complex structures with high F_ident_ to training examples; presence of nucleic acid chains; small total sequence length; large number of unique chains; presence of small molecule ligands; and presence of modified or non-canonical residues in chains. The ability to directly connect features of the underlying structures to Boltz-2 and OpenFold 3 prediction performance profiles demonstrates the nonrandom nature of failure and success modes and points to specific structural features or motifs present among given clusters that lead to the performance profiles observed (See further discussion in Appendix D). Specific cases arising from cluster analysis are discussed in the subsequent subsections.

### Conventional Homology Filtering has Limitations in Preventing Data Leakage

A conventional method for homology attrition of datasets when evaluating new models is using a minimum F_ident_ cutoff. This involves calculating the F_ident_ for each chain in a structure compared to each chain appearing in the set of all structures in the training set of the model. For each chain in the current structure, the maximum F_ident_ compared to all the unique training set chains is recorded. If any of the chains in the current structure has a F_ident_ less than or equal to the chosen cutoff, the entire structure is considered sufficiently novel from the training examples and is thus included in the test set. If no sufficiently novel chain exists in the structure, it is filtered out to avoid homology leakage. Many models leverage this filtering scheme, such as Boltz-2, which uses a F_ident_ of 0.6, and Alphafold 3, which uses a F_ident_ of 0.4 (Passaro et al., 2025; Abramson et al., 2024). However, our results demonstrate a notable limitation of this methodology that may lead to inequitable inflation of model performances in comparative studies due to *residual* homology leakage not attrited using a minimum-F_ident_ filtering approach.

In our analysis, we identify two clusters during HDB-SCAN clustering (denoted clusters 2 and 3) where all of Boltz-2’s metric averages are significantly greater than OpenFold 3’s (see Figure 4). All structures in these clusters contain one large chain where F_ident_ = 1 and a small chain where F_ident_ is below the filtering cutoff (comparing to all chains in the union of Boltz-2 and OpenFold 3 training sets). The structures to which they have high F_ident_ in the union of the training sets are 7OKG and 7QB2 respectively, which were released after the OpenFold 3 training cutoff date (2021-09-30), but before the Boltz-2 cutoff (2023-06-01). These cases demonstrate a limitation of minimum-F_ident_ attrition where the vast majority of the polymeric components of a structure have already been exposed to the model during training, but the standard filtering approach does not remove this data point due to a single, often small chain with low F_ident_ to training chains. It is expected that if a model has already been exposed to the vast majority of a structure during training, it will generally perform better on future predictions containing the leaked chains. We refer to data points that are not filtered by minimum-F_ident_ attrition but that do result in near-perfect model performance as *residual homology leakage*.

**Figure 4.**
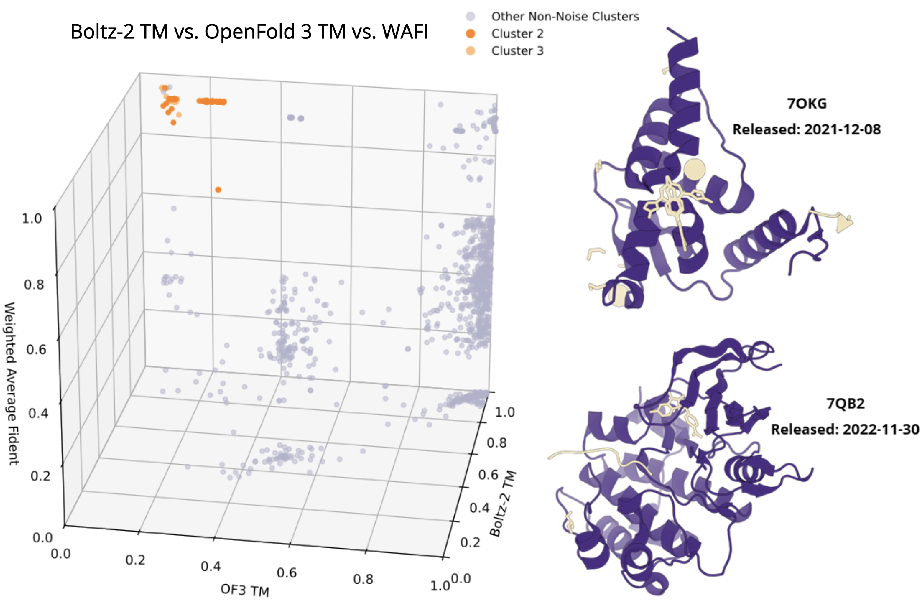
Residual leakage to Boltz-2 training overestimates performance differentials. Comparison of Boltz-2 and OpenFold 3 for TM with Weighted Average F_ident_ (WAFI) plotted along the z-axis (left) and 3D visualizations of the structures to which there is Boltz-2 residual leakage for all structures in clusters 2 and 3 (right).

To prevent residual homology leakage in curating an evaluation set, it is not adequate to include any structure with *one* sufficiently novel chain. This leads to populations of structures which perform abnormally well due to extremely high similarity to training data, but perhaps one short chain with sufficient novelty to avoid attrition. To overcome this limitation of minimum-F_ident_ filtering, we devise an average homology score for a given structure weighted based on each chain’s sequence length (a Weighted Average Fractional Identity score, or WAFI; see the WAFI section in Methods). This prevents instances such as clusters 2 and 3 where a multichain structure may have several large dominating chains with high homology to training examples, but a single small novel chain enables the complex to escape attrition during evaluation set curation.

Altogether, we find that WAFI helps to identify data leakage between homologous structures in training and evaluation sets, providing a more fair comparative evaluation of models with different training cutoffs. However, WAFI is not inherently best suited for all use cases. For example, a campaign aiming to determine the accuracy of small molecule ligand placement would be less concerned with whether a specific filtering scheme results in overinflated estimates of protein folding accuracy; the critical consideration would be the accuracy of the placement of the small molecule. In this case, WAFI may be overly strict, only limiting the test set size without providing additional benefit of gauging model quality for the task of interest. WAFI is primarily advantageous for ensuring biomolecular folding quality is not overestimated in comparisons between models given its improved ability to identify complexes which are highly similar to training examples.

### WAFI Filtering Attenuates Performance Disparities

To better evaluate head-to-head performance, we implement WAFI to further attrite our initial evaluation set to avoid residual homology leakage. This prevents instances where a multichain structure may have several large chains with high homology to training examples, but a single small, relatively novel chain enables the complex to escape attrition during evaluation set curation. Using WAFI in lieu of minimum F_ident_ ≤ 0.6 significantly diminishes the Boltz-2 performance differential over OpenFold 3, decreasing the rate that Boltz-2 scores exceed OpenFold 3 scores by approximately 10 percentage points for all metrics (See Figure 5); the percent of points favouring Boltz-2 now ranges from 51.5% for mean lDDT to 57.4% for mean DockQ. Even so, for lDDT, DockQ, and ilDDT, there are persistent populations with exceedingly poor OpenFold 3 scores; while OpenFold 3 may outperform Boltz-2 on a sizable portion of the evaluation set, when Boltz-2 performs best, the differential is generally more pronounced (while the reverse is not generally true). These results indicate two primary trends: First, WAFI is a more conservative, robust approach to evaluating model generalization to novel examples, and second, using WAFI in place of minimum F_ident_ significantly attenuates if not eliminates any performance differential between Boltz-2 and Open-Fold 3. However, there are persistent populations that OpenFold 3 disproportionately struggles to predict even after correcting with WAFI. We include similar analyses using F_ident_ and WAFI cutoffs at 0.4 in Appendix A within the Supplementary Information.

**Figure 5.**
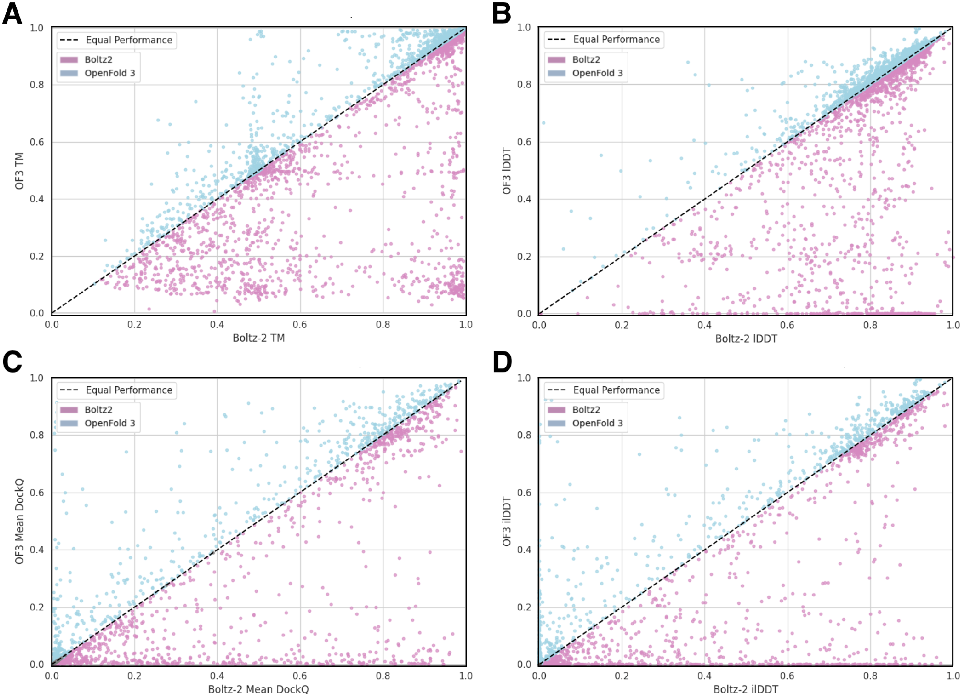
Head-to-head comparison using WAFI in place of F_ident_. (A) TMscores: 55.4% of samples have higher Boltz-2 scores. (B) Mean lDDTs: 51.5% of samples have higher Boltz-2 scores. (C) Mean DockQs: 57.4% of samples have higher Boltz-2 scores. (D) Mean ilDDTs: 55.8% of samples have higher Boltz-2 scores. The provided rates for each plot are the counts of data points below the dashed diagonal, representing the line of equal performance. Inclusion on the plot requires the structure’s overall WAFI *≤* 0.6 compared to the union of Boltz-2 and OpenFold 3 training sets.

### Homology Filtering Strategy and Stringency Significantly Impact Reported Performance Differences

We evaluated the relative performance of Boltz-2 vs. OpenFold 3 under alternative definitions of trainingset homological proximity, again using TM-score, lDDT, ilDDT, and DockQ. For each example *i*, we computed a paired difference *d*_*m,i*_ as defined in Eq. 1:

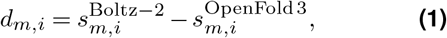

where 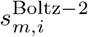 and 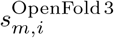 represent the scores for metric *m* from the respective model’s prediction for example *i*. Positive values indicate superior performance of Boltz-2. The dataset was then filtered using two conceptually distinct homology criteria derived from perchain maximum sequence identity to the union of both models’ training sets: 1. Minimum F_ident_, which is the minimum across all chains in the current structure and therefore implements an existential criterion – a complex is retained if there exists at least one chain that is sufficiently novel – and 2. WAFI, a length-weighted average over chains that reflects the overall similarity burden of the complex (see further details on WAFI in Methods). Minimum F_ident_ tends to be a conventional method used in literature for homology attrition of test sets for evaluating new models.

The distinction between minimum F_ident_ and WAFI is fundamental: minimum F_ident_ permits complexes in which most chains may remain highly similar to training data (including potentially to Boltz-2’s training set but not to OpenFold 3’s given Boltz-2’s training cutoff of 2023-06-01 vs. OpenFold 3’s 2021-09-30), provided at least one chain is novel, whereas WAFI penalizes such cases by emphasizing the contribution of larger, potentially non-novel chains. Varying the cutoff (0.6 vs. 0.4) modulates the stringency of these criteria, but under minimum F_ident_, the selection remains intrinsically permissive with respect to partial novelty, whereas WAFI enforces a more global notion of distributional distance from the training sets.

The mean paired-difference ranges and permutationtest results are summarized in Table 2.

**Table 2.**
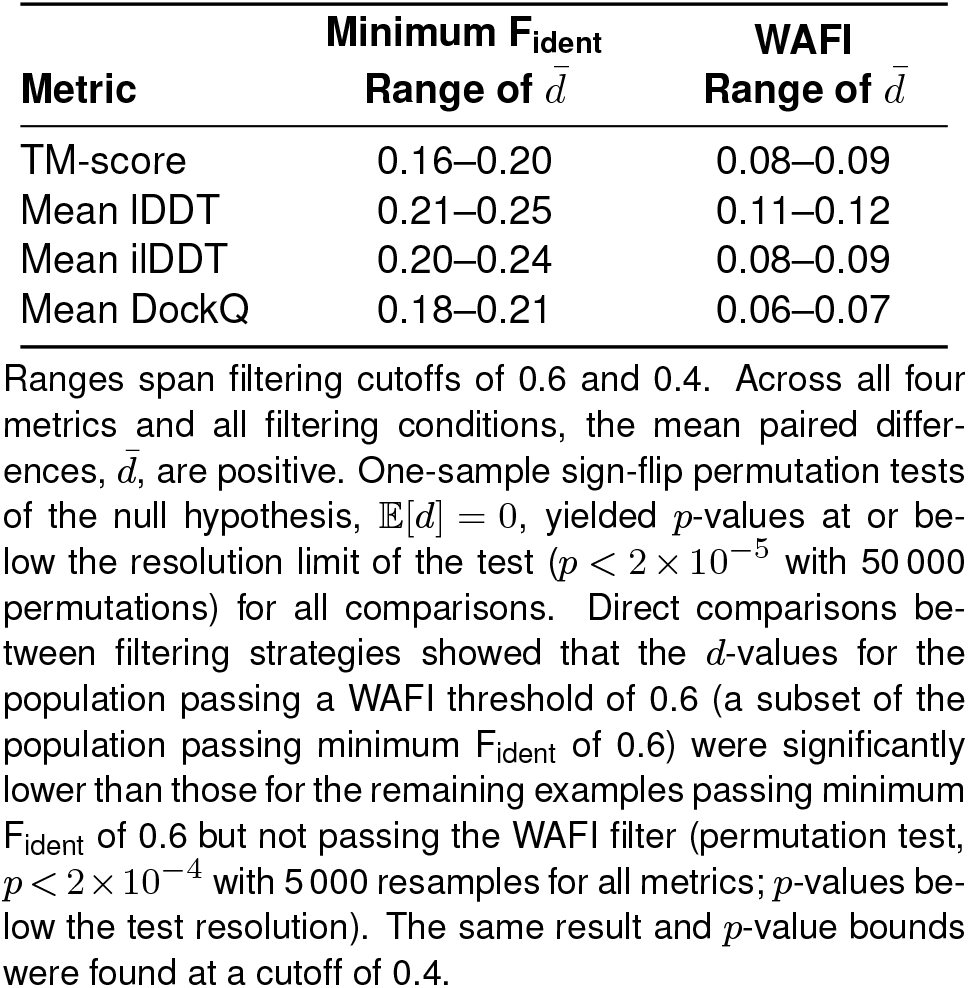
Mean paired performance differences under minimum F_ident_ vs. WAFI homology filters.

|  | **Minimum F<sub>ident</sub>** | **WAFI** |
|---|---|---|
| **Metric** | **Range of $\bar{d}$** | **Range of $\bar{d}$** |
|---|---|---|
| TM-score | 0.16–0.20 | 0.08–0.09 |
| Mean lDDT | 0.21–0.25 | 0.11–0.12 |
| Mean ilDDT | 0.20–0.24 | 0.08–0.09 |
| Mean DockQ | 0.18–0.21 | 0.06–0.07 |

Ranges span filtering cutoffs of 0.6 and 0.4. Across all four metrics and all filtering conditions, the mean paired differences, $\bar{d}$, are positive. One-sample sign-flip permutation tests of the null hypothesis, 𝔼[*d*] = 0, yielded *p*-values at or below the resolution limit of the test (*p* < 2 × 10<sup>−5</sup> with 50 000 permutations) for all comparisons. Direct comparisons between filtering strategies showed that the *d*-values for the population passing a WAFI threshold of 0.6 (a subset of the population passing minimum F<sub>ident</sub> of 0.6) were significantly lower than those for the remaining examples passing minimum F<sub>ident</sub> of 0.6 but not passing the WAFI filter (permutation test, *p* < 2 × 10<sup>−4</sup> with 5 000 resamples for all metrics; *p*-values below the test resolution). The same result and *p*-value bounds were found at a cutoff of 0.4.

These results indicate that Boltz-2’s positive performance differential is robust to the choice of homology filter, but its magnitude depends strongly on the filtering strategy. The systematically lower *d*-values under WAFI suggest that a significant portion of the apparent performance gap under minimum F_ident_ filtering arises from complexes that remain globally similar to Boltz-2 training data despite containing at least one relatively novel chain – precisely the regime WAFI is designed to account for by excluding complexes with high global similarity to training data. Thus, a significant part of Boltz-2’s favourable differential may be attributable to residual homology leakage not excluded by minimum F_ident_ filtering.

The effect of tightening the homology cutoff from a more permissive threshold (0.6) to a more stringent one (0.4) led to another interesting result; when using minimum F_ident_, the *d*-values increased, indicating that OpenFold 3’s performance begins to diverge negatively away from Boltz-2’s with increasing novelty of the most novel chain (*p <* 2 × 10^*−*4^ for all metrics when moving Ranges span filtering cutoffs of 0.6 and 0.4. Across all four metrics and all filtering conditions, the mean paired differences, 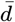, are positive. One-sample sign-flip permutation tests of the null hypothesis, E[*d*] = 0, yielded *p*-values at or below the resolution limit of the test (*p <* 2 *×* 10^*−*5^ with 50 000 permutations) for all comparisons. Direct comparisons between filtering strategies showed that the *d*-values for the population passing a WAFI threshold of 0.6 (a subset of the population passing minimum F_ident_ of 0.6) were significantly lower than those for the remaining examples passing minimum F_ident_ of 0.6 but not passing the WAFI filter (permutation test, *p <* 2 *×* 10^*−*4^ with 5 000 resamples for all metrics; *p*-values below the test resolution). The same result and *p*-value bounds were found at a cutoff of 0.4. from 0.6 to 0.4 with minimum F_ident_). This may suggest greater generalizability to novel sequences for Boltz-2 over OpenFold 3. While a similar statistically significant trend occurs for TM and lDDT (*p <* 2 *×* 10^*−*4^) under WAFI, the change in ilDDT and DockQ are not significant (*p* = 0.082 and *p* = 0.13 respectively). Thus, primarily intrachain/global placement metrics generally diverge in quality with increasing sequence novelty under WAFI, though interchain interactions do not diverge significantly, suggesting Boltz-2’s increased performance differential on highly novel structures is mainly limited to high-quality intrachain predictions as opposed to strong interface predictions.

Taken together, these results indicate that while Boltz-2 demonstrates some advantage over OpenFold 3 across multiple structural metrics, the magnitude of this advantage is substantially inflated under minimum F_ident_-based filtering and becomes more moderate under WAFI-based filtering. This pattern is consistent with the hypothesis that minimum F_ident_ admits complexes with residual training-set similarity that disproportionately benefit Boltz-2 (given Boltz-2’s later training date cutoff than OpenFold 3’s), whereas WAFI more effectively suppresses such cases. Importantly, these findings demonstrate that conclusions about relative model performance are not invariant to the choice of homology filter, and that stricter, globally defined novelty criteria yield more conservative, robust estimates of model generalization.

### Nucleic Acid Complexes Retain Model-Dependent Differences After WAFI Filtering

After WAFI filtering, Boltz-2 retains an advantage over OpenFold 3 for global and local structural metrics in structures with at least one nucleic acid (DNA or RNA) chain, whereas its advantage is smaller for interface metrics (Figure 6). This distinction is reflected in the percent of data points with higher Boltz-2 scores across the filtering conditions, as summarized in Table 3.

**Table 3.** Percent of Data Points with Higher Boltz-2 Score.

| **Condition** | **TM** | **lDDT** | **ilDDT** | **DockQ** |
|---|---|---|---|---|
| F<sub>ident</sub> ≤ 0.6 | 62.5 | 59.5 | 65.8 | 66.9 |
| Contains NA | 79.9 | 77.3 | 69.4 | 71.8 |
| WAFI ≤ 0.6 | 55.4 | 51.5 | 55.8 | 57.4 |
| Contains NA | 78.1 | 74.4 | 58.5 | 64.8 |
| F<sub>ident</sub> ≤ 0.4 | 67.8 | 65.1 | 69.1 | 69.9 |
| WAFI ≤ 0.4 | 59.1 | 54.6 | 55.0 | 57.0 |

The percent of data points for a given filtering condition where the Boltz-2 score is higher than the OpenFold 3 score for the same structure. Dividers delineate a grouping of sequential filters; for example, in the first cell, a filter of F<sub>ident</sub> ≤ 0.6 is first applied, then the requirement that the structure contains a nucleic acid (abbreviated as "NA") is applied. In the next cell, the previous filters are no longer applied, and a new WAFI ≤ 0.6 filter is applied.

**Figure 6.**
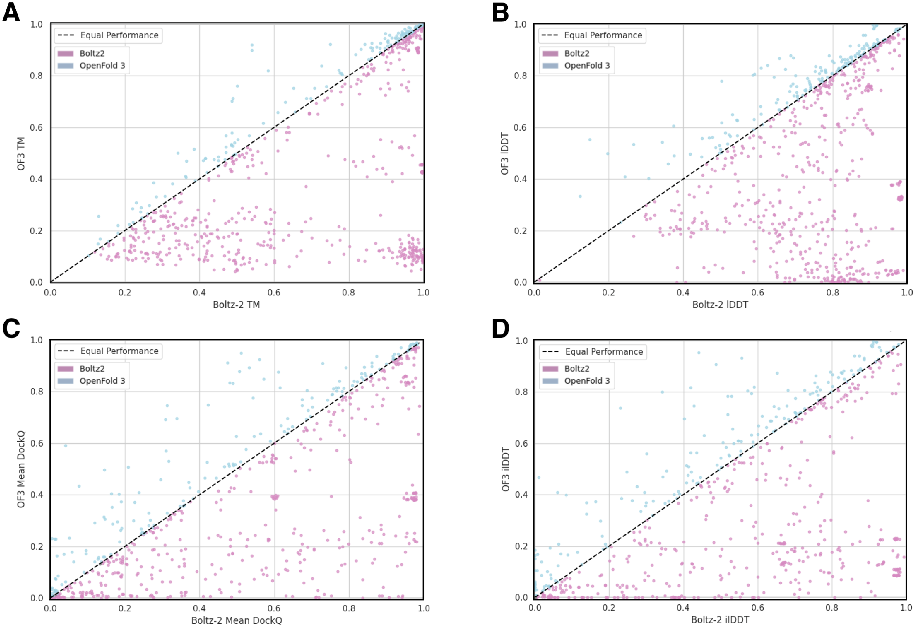
Head-to-head comparison for structures with nucleic acids. General comparison of Boltz-2 vs. OpenFold 3 for structures containing *≥* 1 nucleic acid chain. (A) TM-scores: 79.9% of samples have higher Boltz-2 scores. (B) Mean lDDTs: 77.3% of samples have higher Boltz-2 scores. (C) Mean DockQs: 71.8% of samples have higher Boltz-2 scores. (D) Mean ilDDTs: 69.4% of samples have higher Boltz-2 scores. The provided rates for each plot are the counts of data points below the dashed diagonal, representing the line of equal performance. Inclusion on the plot requires *≥* 1 chain in a structure has F_ident_ *≤* 0.6 compared to the union of Boltz-2 and OpenFold 3 training sets.

The percent of data points for a given filtering condition where the Boltz-2 score is higher than the OpenFold 3 score for the same structure. Dividers delineate a grouping of sequential filters; for example, in the first cell, a filter of F_ident_ *≤* 0.6 is first applied, then the requirement that the structure contains a nucleic acid (abbreviated as “NA”) is applied. In the next cell, the previous filters are no longer applied, and a new WAFI *≤* 0.6 filter is applied.

The substantial increase in data points with higher metrics for Boltz-2 for TM and lDDT (but minimal change for DockQ and ilDDT) may indicate that Boltz-2 performs especially well on global structure and local interaction preservation for structures containing nucleic acids, but that both models generally struggle with predicting interaction interfaces with nucleic acid chains. This is further supported by a visual shift towards the bottom left in the DockQ/ilDDT nucleic acid plots as opposed to the general comparison figures.

When performing the analysis again using WAFI in lieu of minimum F_ident_ *≤* 0.6 (Figure 7), there is once more a decrease in the percent of data points scoring better for Boltz-2 for all metrics, though the decrease is only minor for TM-score and mean lDDT specifically – unlike for the general analysis. This may indicate a more *bona fide* performance differential for Boltz-2 over OpenFold 3 specifically for structures with DNA or RNA given that the rates of better scores for Boltz-2 are relatively preserved despite now applying a more conservative homology filter. This would be an indication of Boltz-2 better generalizing in terms of solving local substructures and global placements for structures containing nucleic acids. This corroborates results described in Passaro et al., 2025, where Boltz-2 is consistently second only to AlphaFold 3 in performance for structures with DNA or RNA, even surpassing AlphaFold 3 performance for rates of high-quality protein-RNA DockQ scores (Passaro et al., 2025). However, in our evaluation set, DockQ and ilDDT scores do see significant drops in the percent of points scoring better for Boltz-2; thus, Boltz-2’s generalization differential is less pronounced in terms of its ability to correctly identify interfaces with nucleic acids.

**Figure 7.**
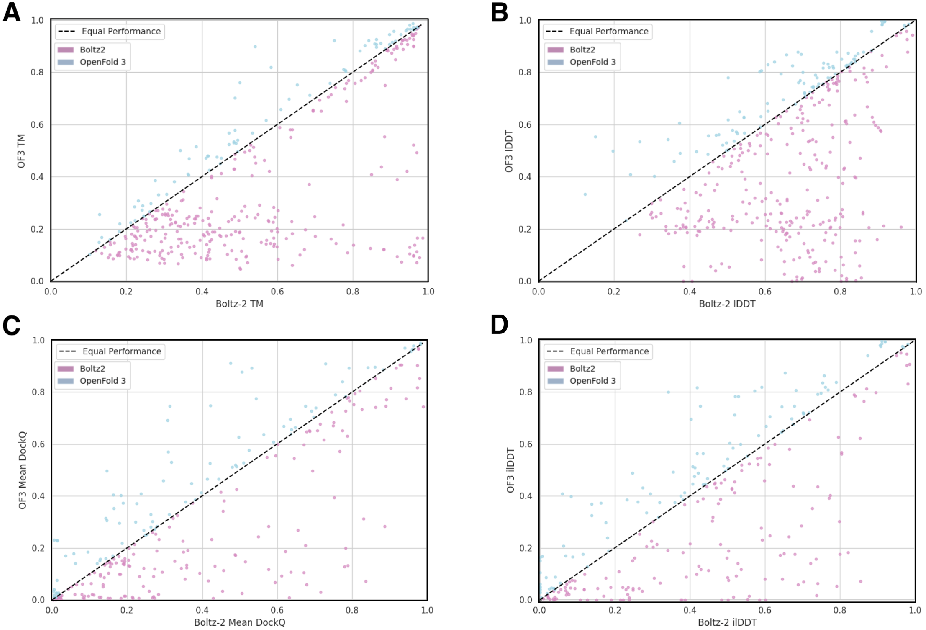
Head-to-head comparison for structures with nucleic acids using WAFI. General comparison of Boltz-2 vs. OpenFold 3 using WAFI for structures containing *≥* 1 nucleic acid chain. (A) TM-scores: 78.1% of samples have higher Boltz-2 scores. (B) Mean lDDTs: 74.4% of samples have higher Boltz-2 scores. (C) Mean DockQs: 64.8% of samples have higher Boltz-2 scores. (D) Mean ilDDTs: 58.5% of samples have higher Boltz-2 scores. The provided rates for each plot are the counts of data points below the dashed diagonal, representing the line of equal performance. Inclusion on the plot requires the structure’s overall WAFI *≤* 0.6 compared to the union of Boltz-2 and OpenFold 3 training sets.

### Homodimer Prediction is Multimodal With Three Primary Failure Modes

Another cluster of particular interest was cluster 5 which had especially poor DockQs for both models. Every structure in cluster 5, including 8UF3 pictured below (see Figure 8), is a homodimer (i.e. two repeated units interacting). Beyond being composed of homodimers, these structures contain no additional modifications or other apparent features that might cause model underperformance. Cluster 5 does not contain all homodimers in the evaluation set. Thus, we further investigated model performance across all homodimers to better understand the divergence in performance from other types of structures.

**Figure 8.**
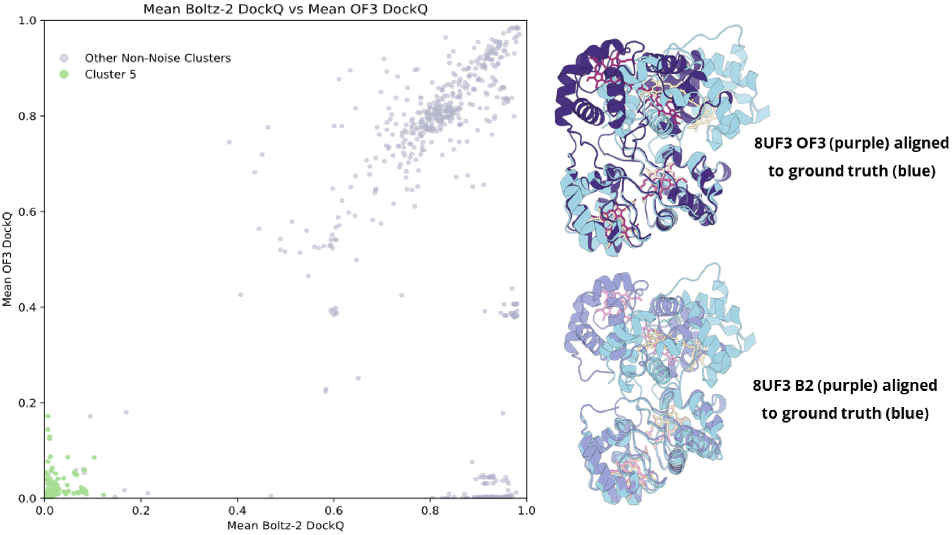
Clusters of homodimers perform particularly poorly. Cluster 5 has poor performance across average DockQ for both Boltz-2 and OpenFold 3 (left). 8UF3 is a structure in this cluster which both OpenFold 3 and Boltz-2 predict poorly.

Plotting metric histograms for all homodimers revealed a sharp multimodality in performance, best illustrated with TM-score (Figure 9). In comparison, heterodimers, monomers and higher-chain-count multimers did not exhibit the same multimodality. Higher order homomultimers exhibit some biomodality as well, however, the bimodality is more gradual and lacking the distinct peak observed in homodimers.

**Figure 9.**
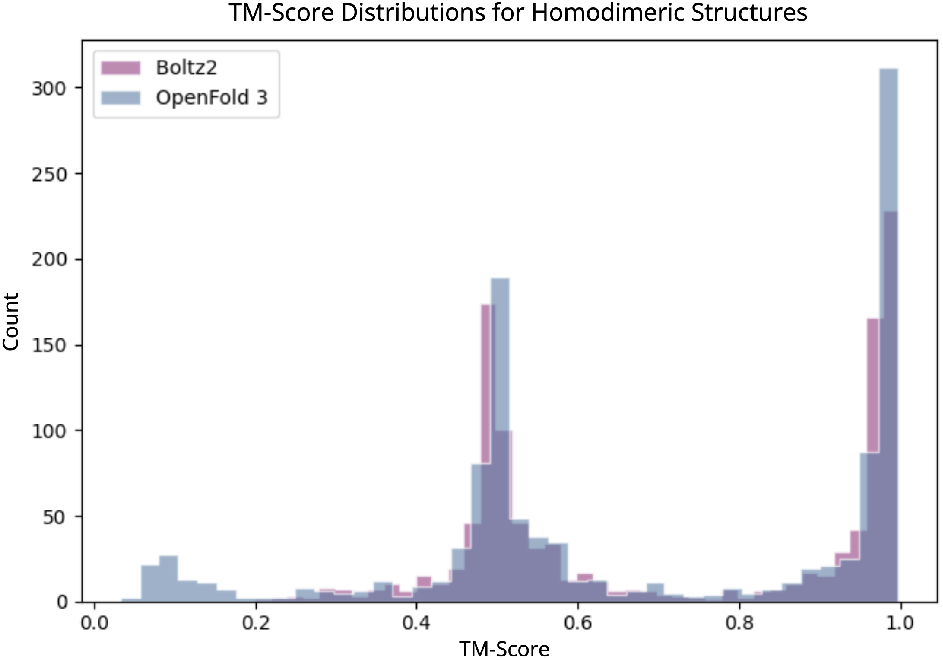
Homodimers show multimodal distributions in TM-score across both models.

To investigate the cause of this sharp multimodality in homodimers, we examined structures which had TMscores of 0.45-0.55 across both model predictions, as well as those which had 0.45-0.55 TM-score for one model and >0.95 for the other. This revealed 3 primary failure modes. The first distinct failure mode is defined by the model predicting the correct interfacial region between the monomers, but the wrong orientation (i.e. one of the monomers might be rotated about the axis normal to the interface; see Figure 10A). Homodimers can show either symmetric or asymmetric binding at the interface. Multiple orientations at the same interface are in some cases plausible, which can lead to model misprediction (Swapna et al., 2012). The second mode of failure is defined by misprediction of the interaction interface (i.e. predicting an incorrect binding site; see Figure 10B). The third failure mode occurs when the ground truth structure differs significantly from the biological assembly (Figure 10C). The ground truth structure published to the PDB is the “Asymmetrical Unit” (ASU), the smallest repeated unit during crystallography. The ASU is not always the same as the biological assembly. In Figure 10C, the ground truth ASU has only a single residue pair closer than 5Å and does not present a reasonable interface. In this case, the ground truth is not a reliable structure for the underlying interaction, and the poor performance simply reflects the unreliable ground truth (as opposed to true model failure).

**Figure 10.**
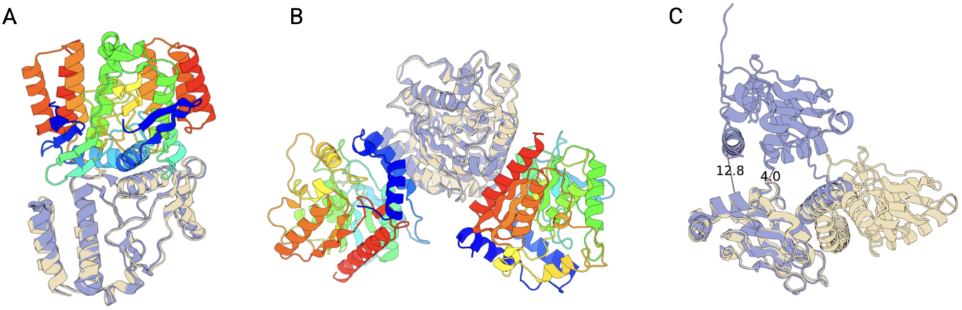
Three examples highlighting distinct modes of homodimeric structure prediction failure. (A) OpenFold 3 prediction (purple and rainbow) overlaid onto ground truth structure (beige and rainbow) for structure 8YV4. The bottom chain is well predicted when aligned; the other is predicted at the correct interface but rotated ~180° (notice the misalignment of chain colours, where blue represents the N terminus and red the C terminus). The OpenFold 3 prediction had TM-score 0.55, while Boltz-2 scored 0.98 (indicating a lack of rotation, i.e. correct orientation at the interface). (B) Boltz-2 prediction (purple and rainbow) overlaid onto ground truth structure (beige and rainbow) for the protein 8YQP. The upper chain is well predicted when aligned, though the other chain is predicted at the incorrect interface as highlighted by the rainbow coloration. Boltz-2 prediction received TM score 0.49, OpenFold 3 prediction scored 0.97 (indicating correct binding site for OpenFold 3, but incorrect site for Boltz-2). (C) OpenFold 3 prediction (beige) overlaid onto ground truth structure (purple) for the protein 8JSK. The ground truth has nearest residue pairs of 4.0 and 12.8 Å. OpenFold 3 and Boltz-2 generate similar structures both with TM-score 0.47. The ground truth structure is more a result of the crystallographic asymmetric unit than the true biological complex, thus leading to reduced performance.

Binding sites are discrete in the sense that there are specific regions of high probability for binding, and others with very low probability. Thus, for cases where multiple physically separate binding regions have similar probability, a model may predict a different interface than the one represented in the ground truth structure. This results in poor alignment and by extension poor structural metrics. Due to the discrete nature of binding sites, we may expect to see multiple Gaussian modes: one near metric scores of 1.0 and others at lower scores. This is especially apparent in TM-score; other metrics such as DockQ show more dispersed points for the same failure modes, but still feature multimodal character.

Compared to their homodimeric counterparts, heterodimer populations in our evaluation set have less stark multimodality. This observation is corroborated by past works, which observe ‘all or none’ behaviour in terms of homomeric interface quality – i.e. that either all interfaces have poor predictions for homooligomeric assemblies, or that all interfaces are predicted well, leading to lower-spread, distinct modes of performances for homomers (Zhu et al., 2023). As for heteromers, each interface in the assembly is structurally unique, so instead of ‘all or none’ behaviour, there is a greater spread of performances. Generally, even modern attention-based models struggle to predict large multimeric assemblies with complex symmetries, and nearly all homomeric assemblies possess a form of internal symmetry, presenting a persisting shortcoming of leading models for high-quality homomeric structure prediction (Kshirsagar et al., 2025; Jeppesen and André, 2023; Zhu et al., 2023).

### OpenFold 3 Oracle Ligands Are More Physically Realistic on Average than Boltz-2’s

To run analysis beyond the structural scoring metrics (DockQ, lDDT, TM-score), we applied PoseBusters to evaluate the physical validity of predicted smallmolecule ligand poses (Buttenschoen et al., 2024). PoseBusters applies 17 checks organized into three tiers:

- Tier 1 assesses chemical validity (sanitization, valence, connectivity, and the absence of radicals)
- Tier 2 assesses internal geometry (bond lengths, bond angles, ring flatness, internal steric clashes, and strain energy)
- Tier 3 assesses protein-ligand context (maximum distance to protein, minimum contact distance, and volume overlap)

A structure is considered a full pass only if it passes all 17 PoseBusters checks across all three tiers. The following analysis focuses on the 1334 structures for which both Boltz-2 and OpenFold 3 produced outputs containing small molecule ligands.

Overall, Boltz-2’s base setup single-structure output achieved a pass rate of 44.4%, while OpenFold 3’s pipeline-recommended sample achieved 35.6% and OpenFold 3’s retrospective most realistic sample (based on PoseBusters checks), defined as the highestpassing sample per structure across all five outputs, achieved 54.2% (Figure 11). OpenFold 3’s recommended sample underperforms Boltz-2 by 8.8 percent, while its most realistic sample outperforms Boltz-2 by 9.8 percent. In essence, the pipeline’s sample selection does not reliably identify the most realistic ligand prediction, and the gap between what the model delivers and what is achievable in principle is 18.6 percent. Investigating these results based on their tier reveals where failures occur. Tier 1 pass rates are nearly perfect for both models (Boltz-2: 99.8%; OpenFold 3 best: 99.8%), indicating that both reliably produce chemically valid small-molecule structures. Tier 2, which evaluates internal ligand geometry, is where the most significant failures occur: only 51.1% of Boltz-2 predictions and 58.0% of OpenFold 3’s most realistic sample predictions pass, meaning roughly half of all predicted poses contain at least one geometric violation at the level of bond lengths, angles, ring planarity, or internal strain energy. Tier 3 pass rates are higher for both models (Boltz-2: 70.4%; OpenFold 3 best: 71.4%), suggesting that the spatial relationship between the ligand and the binding pocket is better preserved than the internal geometry of the ligand itself. This pattern is shown across both models: Tier 2 internal geometry is the primary failure mode, rather than chemical validity or binding site placement. Because OpenFold 3’s standard settings generate five output samples per structure, we were able to analyze variability in PoseBusters outcomes across samples. Of the total OpenFold 3 structures evaluated, 42.9% showed variability across samples, meaning the most realistic sample passed PoseBusters while at least one other sample failed, while the remaining showed no variability, with all samples either passing or failing uniformly. In 35.6% of OpenFold 3 structures, the recommended sample was valid, while in 18.6%, the recommended structure was invalid, and a valid sample existed but was not selected. Finally, in 45.8% no sample passed regardless of selection. We interpret this as evidence that OpenFold 3’s confidence-based sample ranking is not well-calibrated to PoseBusters criteria, and that the 18.6 percentage point oracle gap reflects a selection discrepancy rather than a lacking prediction quality on small-molecules compared to Boltz-2. In fact, if better calibrated, OpenFold 3 may outperform Boltz-2 on small molecule ligand realism.

**Figure 11.**
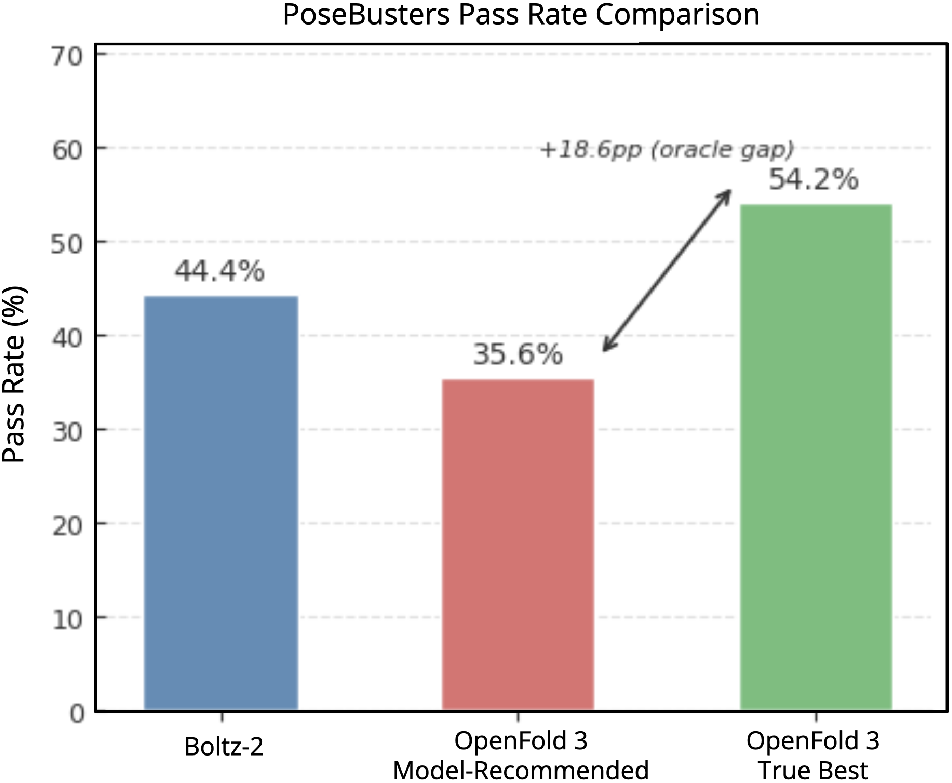
OpenFold 3’s most realistic prediction outperforms Boltz-2’s, though OpenFold 3 often fails to select its most realistic prediction. Bar chart of PoseBusters pass rates for Boltz-2’s oracle, OpenFold 3’s oracle, and OpenFold 3’s most realistic structure (based on PoseBusters checks). Boltz-2’s oracle is more realistic on average than what OpenFold 3 selects as its oracle, though OpenFold 3’s top structure of its five outputs often outperforms Boltz-2’s oracle prediction on PoseBusters pass rate.

### Localization Partitions Reveal Distinct Model Failure Profiles

To explore model performance on complexes present in different subcellular environments, we partitioned data based on subcellular localization identifiers from UniProt annotations. Here, we focus on a simple partition between soluble, membrane and secreted proteins. Figure 12 shows violin plots of DockQ, TM-score, lDDT, and ilDDT for both models stratified by subcellular localization. Performance degrades as proteins move away from the soluble environment that dominates the training data for both models. For Boltz-2, DockQ is strongly top-heavy, TM-score is tightly concentrated near 1.0, and both lDDT and ilDDT are high and narrow. This is consistent with soluble proteins dominating PDB training data and with their folding physics (hydrophobic core burial, hydrogen bond networks in aqueous solution) being well-represented in training (Agarwal and Mc-Shan, 2024). OpenFold 3 shows its characteristic pattern: lDDT and TM have multimodal distributions representing some examples that perform very well and others that are highly inaccurate compared to ground truth. Interface metrics, however, have high failure rates, reflecting interface assembly failures even when individual chains are locally correct. This lDDT-high/DockQ-low dissociation is OpenFold 3’s defining weakness across much of the benchmark.

**Figure 12.**
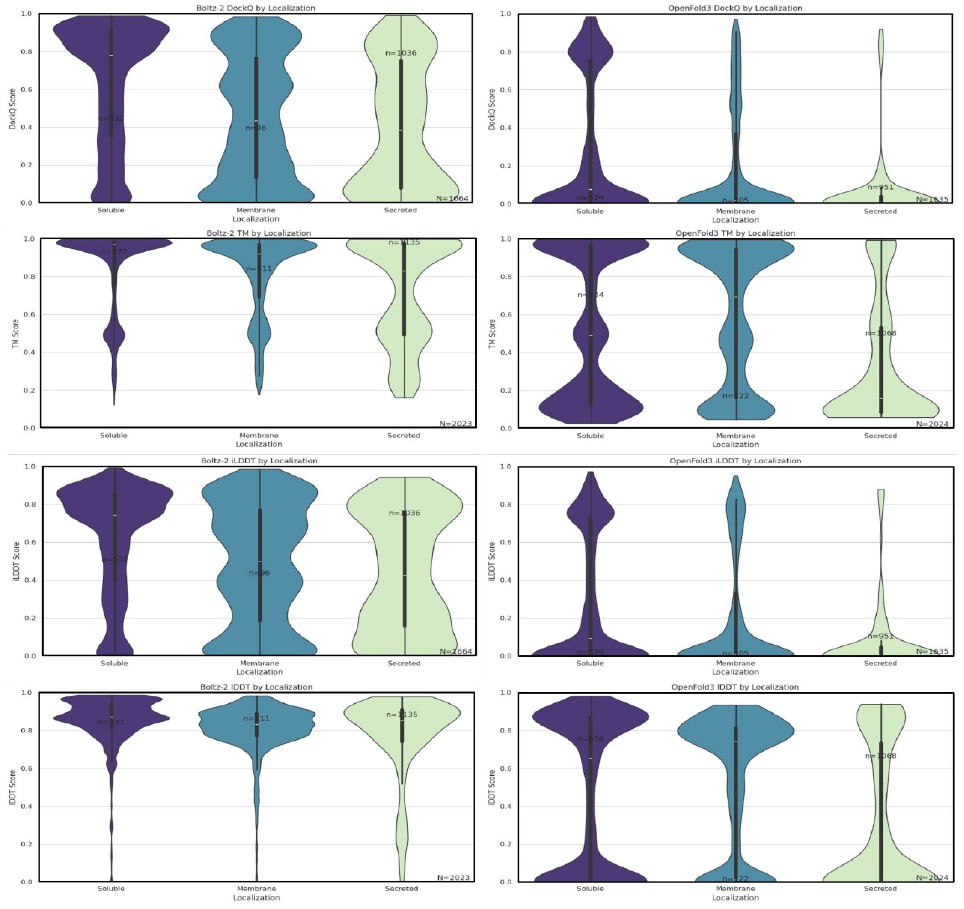
Model performances by subcellular localization. Violin plots of DockQ, TM-score, ilDDT, and lDDT for Boltz-2 (left) and OpenFold 3 (right) stratified by subcellular localization. Both models have best performance on soluble proteins. Membrane proteins show a multimodal Boltz-2 DockQ distribution reflecting partial success on well-represented folds, while OpenFold 3 collapses near zero, consistent with sparse training data and the lipid bilayer being invisible to both models. Secreted proteins show the most severe failures across all metrics, with OpenFold 3 lDDT additionally collapsing.

We expected both models to struggle most with membrane proteins due to their lack of representation in the PDB. Boltz-2 shows a multimodal DockQ distribution for membrane proteins, with one population approaching 1.0 and one near 0. This is a key observation: Boltz-2 is not uniformly worse on membrane proteins but is succeeding on some and failing on others. OpenFold 3’s membrane DockQ is compressed almost entirely to 0, and lDDT and ilDDT are additionally lower than for soluble proteins, indicating that OpenFold 3 loses local geometry quality and especially interface quality for membrane proteins.

We hypothesize three factors that explain this pattern. First, membrane proteins represent less than 3% of PDB structures, meaning both models have been trained on far fewer examples of this class (Agarwal and McShan, 2024). Second, the lipid bilayer physically constrains membrane protein conformation through a hydrophobic core flanked by charged headgroup regions, yet neither model knows the bilayer exists during prediction (Hegedus et al., 2022; Li et al., 2026). Third, a substantial fraction of membrane proteins carry posttranslational modifications including disulfide bonds and N-linked glycosylation that are critical for maintaining topology and structure, none of which are modeled during prediction (Desai et al., 2023).

Secreted proteins show the most severe failures across all metrics for both models, and the failure pattern is qualitatively different from the membrane protein case. For Boltz-2, DockQ is broadly distributed across the full range of values. For OpenFold 3, DockQ and ilDDT collapse toward zero, and even lDDT has a much higher rate of collapse toward zero, a failure mode not observed for soluble or membrane proteins. This failure mode indicates that OpenFold 3 fails at local chain geometry itself, not just interface assembly, for secreted proteins. We hypothesize again that this is due to a mix of underrepresentation in the training data of both models paired with common post-translational modifications on secreted proteins that might not be represented in crystal structures.

### Antibody-Antigen Complexes Remain Challenging for Both Models

We curated a small subset of 57 Ab-Ag pair structures to evaluate model performance on this crucial modelling task. Figure 13 shows paired violin plots of DockQ, TMscore, ilDDT, and lDDT for both models evaluated on the partition. Boltz-2 average scores are higher across all metrics, though both models perform considerably worse on this class than on general protein complexes.

**Figure 13.**
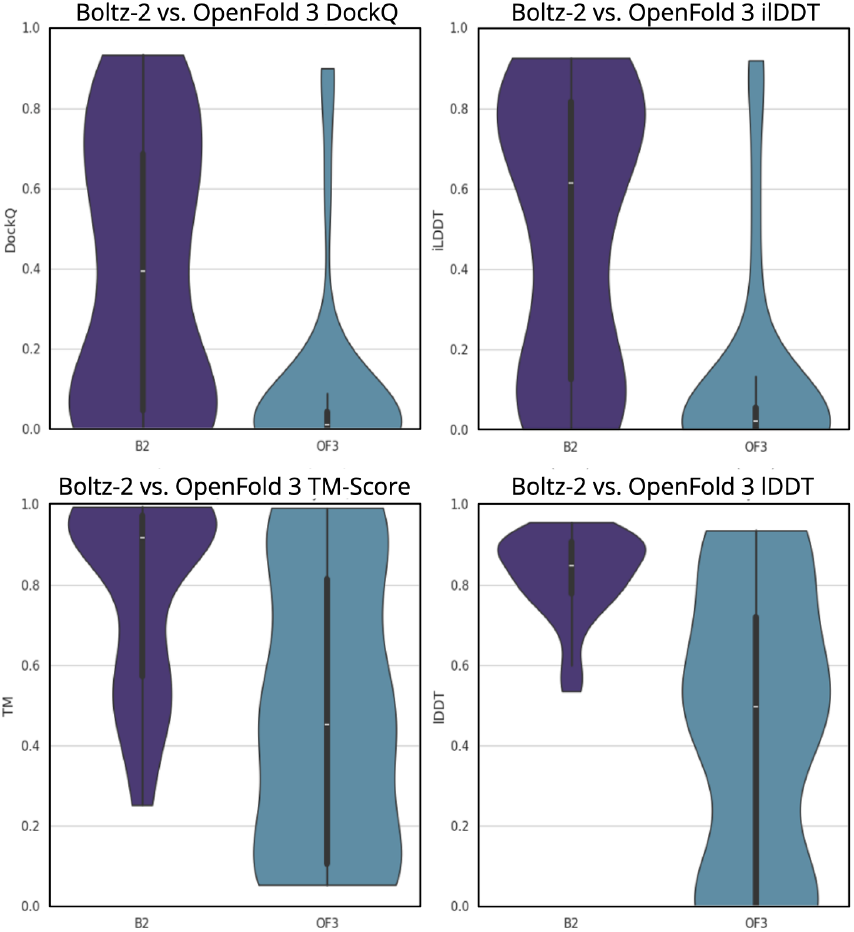
Boltz-2 has higher average metrics on Ab-Ag pairs. Violin plots of DockQ, TM, iLDDT, and IDDT for Boltz-2 and OpenFold 3 on antibody-antigen complexes. Boltz-2 metric averages are higher than OpenFold 3, with OpenFold 3 collapsing near zero on DockQ and iLDDT while Boltz-2 maintains a broad distribution. The struggle generally with interface metrics is consistent with the known challenge of predicting CDR-H3, the most structurally diverse antibody loop, which lacks evolutionary conservation which both models rely on.

OpenFold 3’s DockQ and ilDDT distributions are compressed almost entirely toward zero, representing an exceedingly high rate of interface prediction failure. Boltz-2 maintains a broadly distributed DockQ with a median of approximately 0.4. Both models show broadly spread TM-score distributions, with Boltz-2’s median score around 0.9 and OpenFold 3’s around 0.45, indicating difficulty recovering the correct global fold arrangement, not just interface quality. The lDDT plot is the most diagnostic: Boltz-2’s distribution is narrow and concentrated near 0.87, indicating reliable recovery of intra-chain local geometry, while OpenFold 3 is broadly spread from 0.2 to 1.0 with a median around 0.5, indicating substantial intra-chain geometry failures. The struggle for both models with interface metrics may reflect the inherent complexity and lack of evolutionary preservation among the highly variable CDR loops of the antibody, especially CDR-H3.

### OpenFold 3 Experiences Failed Predictions for Structures with Modified/Non-Canonical Residues

We noticed during analysis that OpenFold 3 struggles with structures containing modified residues in particular. Specifically, these structures have predictions which are fundamentally algorithmic failures (see Figure 14). Any RMSD > 10 Å, or *log*_10_RMSD > 1.0, is generally considered a poor prediction. Viewing the predictions for the population of examples with modified or noncanonical residues, the vast majority of OpenFold 3 predictions have *log*_10_RMSD > 1.0, while only 30% of Boltz-2 structures have *log*_10_RMSD > 1.0.

**Figure 14.**
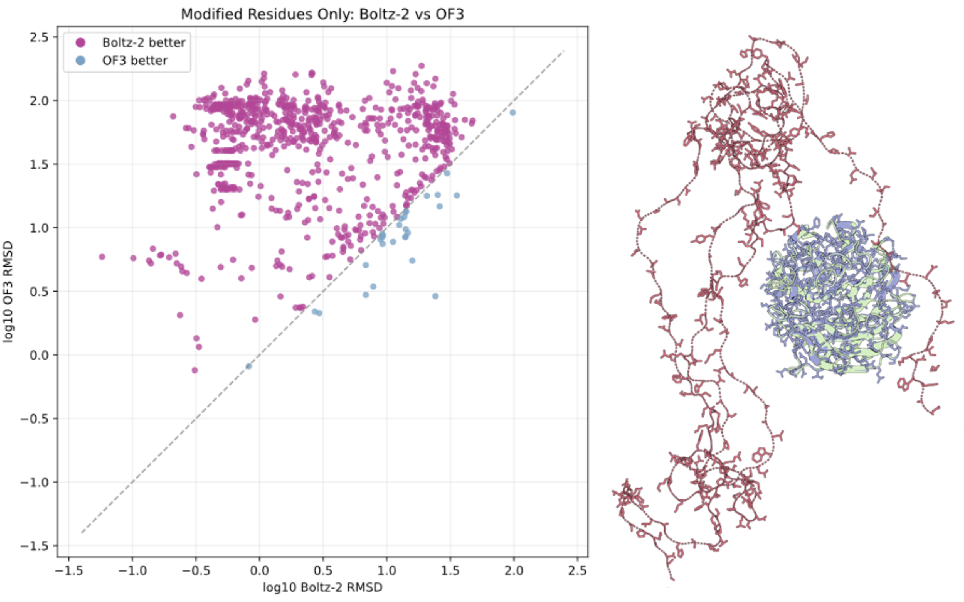
Head-to-head performance on structures containing modified residues. With structures containing modified residues, OpenFold 3 performs significantly worse than Boltz-2, sometimes having RMSD from 10-200Å (left side). When investigating a specific structure (PDB ID: 9UK3) we notice that OpenFold 3 predictions without modifications (green structure) have low RMSD compared to the ground-truth (purple structure) while OpenFold 3 predictions allowing modified residues (red structure) lead to a highly erroneous structure with poor RMSD and many impossible bond lengths.

It is notable that a large portion of Boltz-2 structures similarly have poor prediction quality for structures with modified/non-canonical residues, however, manually inspecting structures, Boltz-2 is able to produce proper secondary and tertiary structures, even if the overall folds are not highly accurate compared to the ground truth. However, manual inspection of OpenFold 3 structures demonstrates frequent unrealistic secondary and tertiary motifs in the predictions. In fact, individual residues are often physically separated far beyond physically realistic bond lengths. As a relatively recently released model, this behaviour is most likely attributable to bugs in the algorithm specifically for structures containing modified residues and will likely be addressed in the near future by the developers if not already by the time of publication of this work.

### Limitations of the Work

While PolyFold provides a robust framework for comparative evaluations of leading biomolecular structure prediction models, several limitations of this analysis warrant discussion.

First, our work is constrained to a pairwise evaluation of two leading open-use models: Boltz-2 and Open-Fold 3. While PolyFold is designed to be fully extensible, this study does not include other common openuse architectures (e.g., Chai, ESMFold, Protenix) nor AlphaFold 3. Extending head-to-head comparisons to a wider spectrum of algorithms remains an important direction for future work.

Second, Boltz-2 and OpenFold 3 incorporate distillation datasets during training. Because the exact compositions of these distillation sets are opaque and difficult to reconstruct from open source, our data leakage mitigation remains incomplete. Although our homology filtering strategies rigorously purge sequence and structural overlap from primary PDB deposits, unaccounted overlap between our evaluation set and the synthetic structures used in the models’ distillation pools may still introduce latent data leakage.

Third, our evaluation pipeline uses PDB ASUs as the experimental ground truth rather than biological assemblies. We selected ASUs because they represent the only direct, un-modeled crystallographic or X-ray coordinate data deposited by experimenters, whereas biological assemblies rely on heuristic or author-curated predictions of functional states. However, because Open-Fold 3 and Boltz-2 are trained to predict physiological biomolecular complexes, instances where the functional biological assembly differs from the crystal ASU can cause metrics to artificially penalize valid predictions that reflect the native interaction rather than the artificial ASU arrangement.

Finally, the models evaluated operate under different training cutoffs: 2023-06-01 for Boltz-2 and 2021-09-30 for OpenFold 3. This temporal gap gives Boltz-2 an inherent baseline advantage, as it was trained on additional deposited structural data. Retraining Open-Fold 3 to align with Boltz-2’s cutoff was computationally prohibitive and unnecessary for establishing PolyFold’s evaluation framework; nevertheless, readers should account for this training window asymmetry when interpreting absolute performance differences.

## Methods

### Evaluation Metrics

#### RMSD

Root Mean Square Deviation (RMSD) serves as a classical metric for assessing global structural divergence by computing the mean distance between *N* corresponding paired atoms — typically C*α* atoms for proteins or backbone P atoms for nucleic acids — following optimal rigid-body translation and rotation. Given coordinates 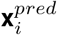 for the predicted model and 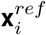 for the reference structure across *i* ∈ {1, 2, …, *N*}, the metric is mathematically expressed as:

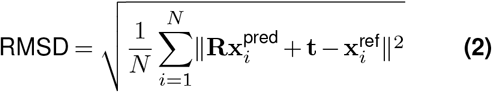

where **R** ∈ SO(3) is the optimal 3 × 3 rotation matrix and **t** ∈ **R**^3^ denotes the translational alignment vector derived via superposition algorithms such as the Kabsch method. RMSD operates on a continuous range of [0, ∞), where 0Å signifies perfect structural superposition, and higher values correspond to decreasing global accuracy. RMSD is inherently scale-dependent and highly sensitive to conformational variations; because distance errors are squared, localized deviations within flexible tails, unstructured loops, or domain hinge rotations can artificially inflate the score despite identical tertiary core folds. Thus, RMSDs are often logscaled, or other global structure metrics such as TMscore are used.

#### TM-score

The Template Modeling score (TM-score) addresses the key limitations of RMSD by evaluating global protein fold architecture through a scale-independent, distanceweighted function that attenuates the impact of large spatial deviations in mobile loops while emphasizing conserved core topology. For a target reference structure of sequence length *L*_target_ and an aligned pair count of *L*_aligned_, TM-score is defined as:

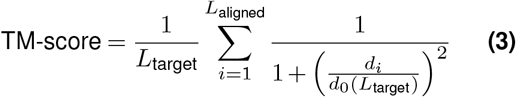

where *d*_*i*_ represents the Euclidean distance between the *i*^th^ aligned pair of C*α* atoms post-superposition (following the same methodology as RMSD for superposing the predicted model to the reference structure), and 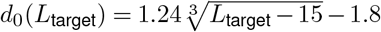 is a dynamic scaling distance that standardizes the sensitivity of the metric relative to average protein dimensions. *d*_0_ assumes *L*_target_ *>* 21, but for short peptides where *L*_target_ ≤ 21, *d*_0_ is set to a lower bound of 0.5Å). The alignment is optimized over coordinate space to maximize the overall summation. The TM-score yields a strictly bounded continuous value in (0, 1], where 1.0 indicates perfect identity. A TM-score threshold of 0.5 or greater serves as a statistically rigorous cutoff signifying that the predicted and reference structures share the same global tertiary fold family (*p <* 0.001), whereas values falling below 0.17 correspond to random structural pairs possessing no meaningful topological relationship.

#### lDDT and ilDDT

The Local Distance Difference Test (lDDT) is a superposition-free metric that measures the local stereochemical correctness and atom-packing environment of a predicted structure by evaluating internal pairwise distances rather than global Cartesian coordinates. For a reference structure, a subset *S* is constructed containing all unique atom pairs (*i, j*) separated by an internal distance 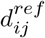 within a local inclusion sphere radius *R*_*cut*_ (usually 15.0Å), omitting immediate sequence neighbours such that |*i* − *j*| *>* 1. The absolute distance error for pair *k* ∈ *S* is computed as 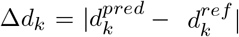. Evaluating these errors across four predefined tolerance thresholds *T* = {0.5Å, 1.0Å, 2.0Å, 4.0Å}, the global lDDT score is defined as:

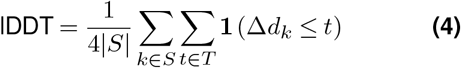

where **1**(·) is an indicator function returning 1 if the distance error remains within tolerance threshold *t* and 0 otherwise. The resulting score ranges continuously between 0 and 1, with 1.0 reflecting absolute local distance preservation. Because lDDT avoids rigid-body coordinate alignment, it is immune to rigid domain movements or inter-domain hinge movements that confound global metrics. For multi-chain structural assemblies, local binding interface preservation can be measured using interface lDDT (ilDDT). In ilDDT, the evaluation set *S*_*interface*_ is restricted strictly to inter-chain atom pairs where atom *i* and atom *j* belong to different chains (Chain(*i*)≠Chain(*j*)) with 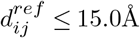. Computing the four-threshold average over *S*_*interface*_ evaluates interprotein packing contacts without interference from intrachain backbone errors.

#### DockQ

DockQ is a continuous quality metric designed to assess multi-chain macromolecular complex predictions by unifying three traditional, discrete Critical Assessment of PRediction of Interactions (CAPRI) metrics into a single continuous score bounded within [0, 1]. Derived from non-linear scaling transformations calibrated to reproduce historical CAPRI quality standards, DockQ is calculated as:

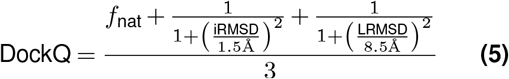

The first subcomponent, the fraction of native contacts (*f*_*nat*_), measures interface recovery by evaluating interchain heavy atom pairs within 5.0Å of each other. Expressed as:

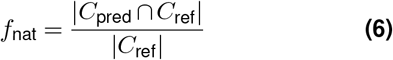

where *C*_*ref*_ and *C*_*pred*_ represent the native and predicted contact sets respectively, *f*_*nat*_ quantifies the proportion of correct interface interactions. The second subcomponent, interface RMSD (iRMSD), quantifies local interface backbone geometry; it is computed as the RMSD over backbone atoms (C*α*, C, N, and O) belonging to interface residues (defined as residues possessing any heavy atom within 10.0Å of a neighboring chain in the reference complex) following optimal superposition restricted strictly to these interface backbone atoms. The third subcomponent, ligand RMSD (LRMSD), captures the global relative orientation of the complex by superposing the backbone of the larger stationary partner (receptor) and measuring the resulting backbone RMSD of the unaligned smaller partner (ligand). Based on established CAPRI benchmarks, predictions evaluated via DockQ are categorized as incorrect (DockQ *<* 0.23), acceptable quality (0.23 ≤ DockQ *<* 0.49), medium quality (0.49 ≤ DockQ *<* 0.80), or high quality (DockQ ≥ 0.80).

### Evaluation Set Construction

Information leakage is an increasingly important consideration in evaluating biomolecular structure prediction algorithms (Joeres et al., 2025; Teufel et al., 2023). When assembling an evaluation set for these models, it is not sufficient to simply exclude data points present in the training or validation set of the algorithm under investigation (as is the case in other machine learning domains); it is necessary to additionally compare sequence homology against the training set to ensure realistic reporting of model performance on out-ofdistribution data. This process entails applying homology reduction to the evaluation set. In the context of this work, homology reduction is attriting the evaluation set by removing structures where every chain in the structure has a homology similarity score which exceeds a pre-defined similarity threshold. The similarity score is calculated by comparing a chain to all chains occurring in the training set of the model (or models) under evaluation and reporting the maximum fractional identity score, F_ident_, from 0 to 1 (1 being perfect alignment). This filtering framework is summarized as follows (Equations 7, 8, and 9):

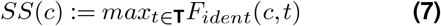

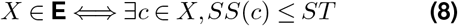

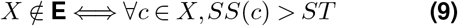

Where *SS*(*c*) is the similarity score for chain *c*, **T** is the set of chains appearing in the training set of the model under evaluation, *F*_*ident*_(*c, t*) is the fractional identity score between chain *c* and chain *t* (as calculated by MMseqs2 when running the search command), *X* is an arbitrary structure potentially composed of several chains, **E** is the evaluation set, and *ST* is the similarity threshold. As an example, this homology reduction framework would imply that for a 5-chain structure *X*, if even 4 of 5 chains have high *SS*, the structure will not be attrited because there is at least 1 chain *c* with adequate dissimilarity to all chains *t* ∈ **T** (i.e. the chains in the training set of the model under evaluation).

In relation to evaluating OpenFold 3 and Boltz-2, we assembled a 6063-structure set of post-training examples, **E**. To accomplish this task, we first collected all training examples from OpenFold 3 and Boltz-2: **T**_*OpenF old*3_ and **T**_*Boltz−*2_ respectively. From here, we performed homology reduction to create our evaluation set by comparing all structures after the last validation cutoff date (2023-12-31; see Figure 15) to the union of the sets (as defined in Equation 10 below):

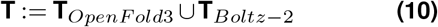

We started by assembling **T**_*OpenF old*3_ then continued to Boltz-2 data in order to assemble **T**. The process for recreating **T** from **T**_*OpenF old*3_ and **T**_*Boltz−*2_ involved filtering structures based on the pipeline described in the Boltz-2 and OpenFold 3 pre-prints. Namely, we:

**Figure 15.**
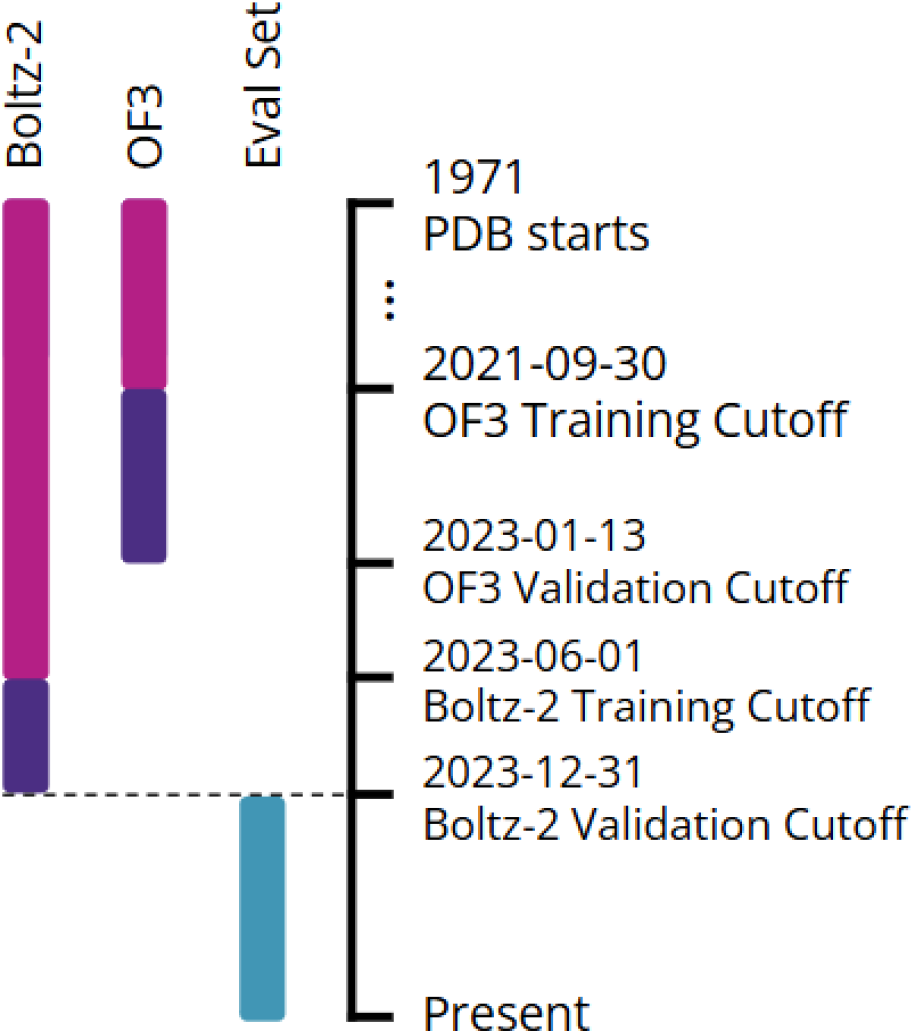
Model key cutoff dates. Timeline detailing Boltz-2 and OpenFold 3 training and validation cutoffs, providing insight into the starting point for assembling the evaluation set described in this work.

- Take all PDB structures up to the latest training cut-off (Boltz-2’s training cutoff of 2023-06-01)
- Remove large structures greater than 7 MB or 5000 total residues (the sum of sequence lengths of all polymer chains)
- Remove structures with resolution worse than 9 Å
- Apply the same filters as AlphaFold 3, namely excluding crystallization aids and other nonbiologically relevant ligands, and filtering out chains with fewer than 4 resolved residues or composed only of unknown residues.

Notably, we did not attempt to reassemble the distillation data additionally used to train Boltz-2 or OpenFold 3; information on this data is sparse, making replication difficult. This is an important limitation to factor in with our analysis, as it may lead to biased information leakage not accounted for in our homology attrition.

After reassembling **T**, we took all structures post-2023-12-31 (i.e. Boltz-2’s validation set cutoff date, the later of the two validation sets) and performed the following filtering operations:

- Homology attrition: For all unique polymer chains in post-2023-12-31 structures, we collected all nucleic acid chains to one FASTA and all protein chains to another. We did the same for all chains in **T**. We then created MMseqs2 databases from the FASTA files and ran two MMseqs2 search commands with a sensitivity of 7.5 (all other parameters left as their default value): one for **T** vs. post-2023-12-31 protein sequences using searchType 1, and one for **T** vs. post-2023-12-31 nucleotide sequences using searchType 3. This generated multiple sequence alignments for all unique polymer chains in post-2023-12-31 structures based on all structures in **T**. All structures which lack a single chain with F_ident_ ≤ 0.6 (i.e. whose *SS* was greater than the cutoff) were removed from the set given they are considered too similar to training examples
- Largely following Boltz-2’s test set assembly, for interfaces, we keep only interfaces where at least one of the two chains is sufficiently distinct, as defined above
- We remove targets with resolution worse than 4.5 Å. If a resolution is not provided, the structure is similarly removed
- Structures greater than 7 MB or longer than 1200 residues in total residue length are removed (to save on computational resources for predictions with large memory requirements)

At this point, we arrive at our defined set **E**. By applying a 0.4 *SS* cutoff to **E** or using WAFI to further filter **E** (using both WAFI ≤ 0.6 or ≤ 0.4), we assembled more stringent subsets of **E** for analysis of the effect of the homology filtering method on relative model performances.

### Data Partitioning of the Evaluation Set

To provide a more specific and informative analysis of **E**, we partitioned our data into various subsets for more granular performance comparisons. Structural metadata (e.g., resolution, experimental method for structure derivation, and assembly composition) were extracted from PDB CIF files, while composition and content features (e.g., small molecule presence, modified residues, etc.) were derived by analyzing CIF and FASTA files as well as scraping information from the PDB records. Functional and biological context, including enzyme classification (EC numbers), subcellular localization, UniProt annotations, and InterPro family assignments, were incorporated via external database mappings. Chain-level and global sequence features, such as sequence identity and weighted average identity, were computed from alignment-based analyses. Structural prediction outputs from Boltz-2 and Open-Fold 3 were linked to each entry, and their performance was evaluated using OpenStructure (version 2.11) to compute global, interface, and ligand-specific accuracy metrics (e.g., RMSD, TM-score, lDDT, DockQ) (Biasini et al., 2013). Additional quality control metrics and error flags were included to track structural validity and evaluation success. For small molecule realism checks, we used PoseBusters (version 0.6.4) (Buttenschoen et al., 2024). Together, these features provide a unified representation of structure, function, and predictive performance, enabling systematic downstream analysis (see Appendix B in the Supplementary Information for detailed definitions of all variables).

### Weighted Average Fractional Identity

To replace the information leakage-prone minimum F_ident_ approach to homology reduction, we conceptualized the Weighted Average Fractional Identity, or WAFI, which we use throughout this work for a more stringent alternative to minimum F_ident_. WAFI is defined as follows (Equation 11):

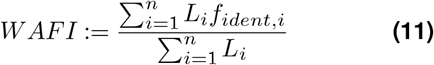

where *n* is the total number of chains in the structure, *L*_*i*_ the sequence length of chain *i*, and *f*_*ident,i*_ the fractional identity of chain *i* to the homologically “closest” training set sequence. It is fundamentally a sequencelength-weighted average fractional identity score.

### PolyFold: A Workflow Automation Tool

We created an integrated workflow automation tool for streamlining input file creation, inference of biomolecular structure prediction algorithms, and evaluation of model outputs (see the GitLab repository here). This package will allow scientists to parallelize thousands of inference commands with only a few lines on the terminal: one to generate PolyFold input files from structure files if required (namely, a FASTA file for sequences, a CSV for ligand SMILES strings / CCD codes, and a JSON for residue modifications and noncanonicals), one to generate input files for the structure prediction algorithm, one to run the job from the workload manager script produced, and two to evaluate structures (one for OpenStructure structural quality evaluation and one for PoseBusters small molecule realism evaluation (if required)).

Our initial release provides compatibility with Boltz-2, OpenFold 3, and AlphaFold 3, though we encourage potential collaborators to contribute in expanding the suite of compatible models by referring to the README in our repository.

### Clustering Examples on Output Metrics

We performed clustering of datapoints based on the main structural performance metrics from both the Boltz-2 and OpenFold 3 outputs for that example. We vectorized each datapoint into 20 dimensions, where each dimension is a Z-score-normalized structural performance metric. The 20 dimensions are as follows: 1. Boltz-2 mean DockQ (averaged across all interfaces in the structure), 2. Boltz-2 mean ilDDT (across all interfaces), 3. Boltz-2 lDDT, 4. Boltz-2 *log*_10_RMSD, 5. Boltz-2 TM-score, 6. Boltz-2 *log*_10_(Mean Ligand RMSD), 7. Boltz-2 mean ligand lDDT, 8. Boltz-2 number of “bad bonds” (based on OpenStructure), 9. Boltz-2 number of “bad angles”, 10. Boltz-2 number of model clashes, 11-20. The same dimensions, but for OpenFold 3 results. We clustered both with this 20-dimension vectorization as well as with a 10-dimensional approach (where we only used dimensions 1-5 and 11-15). We used HDB-SCAN cluster size of 20, which provided an appropriate balance of clustering and non-clustering datapoints. Appendix D in the Supplementary Information investigates how manipulating the cluster size parameter affects the clustering outcomes.

### Effect Size and Multivariate Regression Across and In Clusters

#### Effect Size (Cluster vs. Out-Of-Cluster Differences)

For the clusters obtained from HDBSCAN, we performed an effect size analysis to determine which input variables drive clustering significantly. The input variables we used are: presence of ions, presence of small molecule chains, presence of modified residues, nonpolymer entity count, nucleic acid chain count, protein chain count, number of unique chains, sequence length, and WAFI. Because our inputs are binary, discrete, and continuous, we used different statistical methods which we then adjusted using the Benjamini-Hochberg false discovery rate procedure. Predictors with an adjusted *q*-value below 0.05 were considered statistically significant.

For continuous predictors (such as WAFI), differences between groups were assessed using the two-sided Mann-Whitney U test. Effect sizes were quantified using the rank-biserial correlation, derived from the Mann-Whitney U statistic, to capture the direction and magnitude of distributional differences without assuming normality. For binary predictors (e.g., presence of smallmolecule ligands, ions, or modified residues), associations were evaluated using Fisher’s exact test on 2 × 2 contingency tables. Effect sizes were defined as the difference in proportions between the cluster and the remainder of the dataset. All analyses were performed on the full dataset used for clustering, with effect sizes and significance values computed independently for each cluster-predictor pair. The statistics and other results are detailed in Appendix D within the Supplementary Information.

#### Multivariate Regression (In-Cluster Model Differences)

Following effect size analyses, we performed clusterspecific multivariate regression on the same inputs as above to identify structural and compositional features associated with model performance discrepancies between Boltz-2 and OpenFold 3. For each cluster, the dependent variable was defined as the paired performance gap between Boltz-2 and OpenFold 3, computed for each of the following structure metrics (*m*): Average DockQ, lDDT, TM, iLDDT, and *log*_10_RMSD (Equation 12):

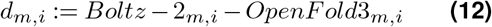

such that positive values indicate superior performance of Boltz-2 on metric *m* for example *i*. Predictor variables were preprocessed prior to running the regression: continuous variables were log-transformed and standardized to zero mean and unit variance (computing a Z-score for each value), while binary variables were encoded as 0 or 1. Ordinary least squares regression was then applied independently for each performance metric within each cluster. To account for potential heteroskedasticity, robust standard errors (HC3) were used for inference. Model coefficients were interpreted as the direction and magnitude of association between each predictor and the relative performance advantage of Boltz-2 over OpenFold 3.

Collinearity among predictors was assessed using variance inflation factors, and model fit was evaluated using *R*^2^, adjusted *R*^2^, and information criteria (AIC, BIC). Residual diagnostics, including residual-fitted plots and Q-Q plots, were used to assess model assumptions. An example of this complete analysis can be seen in Appendix D within the Supplementary Information.

### Statistical Analysis of Paired Differences Using F_ident_ vs. WAFI at 0.4 and 0.6

Per-example structural quality was quantified using four metrics: TM-score, lDDT, ilDDT, and DockQ. For each metric *m* and example *i*, a paired performance differential was defined identically to Equation 12 in the previous section, such that positive values indicate superior performance of Boltz-2 on metric *m*.

#### Homology Filtering Criteria

Using either minimum F_ident_ or WAFI cutoff methods for homology reduction (as described above), both criteria were applied at thresholds of either 0.6 and 0.4 to attrite the evaluation set. For each metric and filtering condition, the corresponding subset of PDB identifiers was extracted and used to compute summary statistics.

#### One-Sample Sign-Flip Permutation Test

To assess whether Boltz-2 significantly outperformed OpenFold 3, we tested the null hypothesis for a given mean paired difference value, 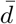 (i.e. E[*d*] = 0) using a one-sample sign-flip permutation test. This nonparametric test is appropriate for paired differences and does not assume normality; it relies only on producing symmetry of the null distribution around zero.

For a given set of observed differences, *d*_*i*_, the test proceeds by randomly assigning each *d*_*i*_ a sign of +1 or −1 with equal probability, generating permuted datasets under the null hypothesis. For each of 50 000 resamples, the mean of the sign-flipped differences was computed. The two-sided *p*-value was estimated as the proportion of permuted means with absolute value greater than or equal to the observed mean, using the standard finite-sample correction (Equation 13):

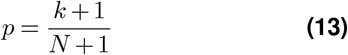

where *k* is the number of extreme permutations and *N* is the number of resamples. This yields a minimum attainable *p*-value (i.e. the maximum resolution of the test) of approximately 2 × 10^*−*5^.

#### Subset vs. Population Permutation Tests

To compare performance differentials between subsets (e.g., WAFI-filtered complexes with a 0.6 cutoff) and their complementary populations, we used a twosample permutation test. Since a WAFI-filtered subset at 0.6 cutoff is a direct subset of the evaluation set, **E**, comparing the mean paired differences for the WAFI subset vs. **E** minus the WAFI subset gives an indication of whether using WAFI in place of minimum F_ident_ results in a significantly different reported performance differential between Boltz-2 and OpenFold 3 for a given performance metric.

For a given subset **S** of size *k* drawn from a total pool **P**, the observed statistic was defined as:

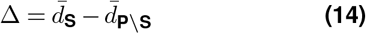

Under the null hypothesis that subset membership is unrelated to performance differential, labels were permuted by repeatedly sampling random subsets of size *k* from **P** without replacement. For each of 5000 resamples, the difference in means was recomputed. Twosided *p*-values were calculated analogously to the onesample test.

This framework was used to:

- Compare each filtered subset to the full evaluation set (i.e. minimum f_ident_ filter at 0.6 cutoff). The filtered subsets were: minimum f_ident_ at 0.4 cutoff, WAFI at 0.6 cutoff, and WAFI at 0.4 cutoff
- Compare the WAFI at 0.4 cutoff subset to the pool, when the pool is redefined as the minimum f_ident_ at 0.4 cutoff set (since WAFI at 0.4 is a direct subset of minimum f_ident_ at 0.4)
- Compare the WAFI at 0.4 cutoff subset to the pool, when the pool is redefined as the WAFI at 0.6 cutoff set (since WAFI at 0.4 is a direct subset of WAFI at 0.6)

#### Implementation Details

All analyses were implemented in Python using numpy and pandas. Random number generation was controlled with a fixed seed to ensure reproducibility. All statistical tests were performed directly on aligned arrays of paired differences indexed by PDB ID. The number of permutations (50 000 for one-sample tests; 5000 for subset comparisons) was chosen to balance computational efficiency with sufficient resolution for small *p*-values. All reported *p*-values are two-sided and reflect the empirical permutation distribution. This nonparametric, permutation-based framework ensures robustness to distributional assumptions and is well-suited to the paired, heterogeneous nature of structural prediction performance data.

### Excluded Ions, Small Molecules, and Crystallization Aids During PoseBusters Analysis

Following a similar approach to AlphaFold 3, we excluded specific molecules during PoseBusters evaluation of small molecule realism. The rationale guiding exclusion is that many molecules do not contribute meaningfully to the performance of a model in terms of small molecule realism, namely crystallization aids and some single-atom ions (which cannot have internal molecular RMSDs, lDDTs, or problematic bond lengths or angles since they are composed of a single atom). Following the conventions of AlphaFold 3, we excluded any molecules listed in Appendix C of our Supplementary Information (Abramson et al., 2024).

## Conclusions

In this work, we developed PolyFold, an end-to-end, reproducible benchmarking framework for evaluating structure prediction models on various biologically relevant folding tasks. Using PolyFold to evaluate 6063 curated structures, a key finding of our work is that benchmarking outcomes are highly sensitive to how structural novelty is defined. Conventional minimumchain fractional identity filtering inflates apparent performance differences due to residual homology leakage. By introducing weighted average fractional identity, we demonstrate that shifting to structure-level homology control significantly affects the conclusions drawn in head-to-head model comparisons. Under conventional novelty filters, Boltz-2 was found to have higher average metrics across multiple biologically important tasks, including antibody-antigen complexes, nucleic acid-containing structures, and structures with modified residues. However, this performance differential is reduced or even eliminated when more stringent homology filtering methods are imposed. This underscores the limitations of conventional evaluation practices when attempting to accurately assess model generalization. Beyond aggregate performance, we show that model success is strongly task-dependent and governed by structured failure modes. Both models continue to struggle with complex interface prediction — particularly in homomultimers and antibody-antigen systems — as well as with small molecule realism and structures containing non-canonical amino acids. These limitations are systematic rather than random, highlighting the importance of task-specific model selection.

Together, these developments position PolyFold as a scalable and extensible benchmarking platform for biomolecular structure prediction. Beyond the comparison presented here, PolyFold provides a reusable infrastructure for the rigorous, transparent, and taskaware evaluation of future models. As structure prediction methods continue to evolve rapidly, standardized benchmark construction, careful control of training-data leakage, and reproducible evaluation protocols will be essential for ensuring that reported improvements reflect real advances in model capability rather than artifacts of benchmark design.

## Supporting information

Evaluation Set Data

## DATA AND CODE AVAILABILITY

The curated evaluation set used in this study, including PDB identifiers, model evaluation metrics, structural input data, and associated metadata, is provided as the supplementary CSV file titled *Evaluation Set Data*. The PolyFold source code used in this study is maintained in the GitLab repository at https://gitlab.com/congruencetx/polyfold. The version used for this study corresponds to Git commit 1285cad1044b7f046af9f9f0c7f49af0fb5e0519. The repository will be made publicly accessible upon publication.

## ACKNOWLEDGEMENTS

This work was initiated in the context of a capstone course in McGill University’s Department of Bioengineering. We thank Congruence Therapeutics for guidance and scientific discussion throughout the project, as well as access to computational resources. We thank Jasmin Coulombe-Huntington for his guidance and constructive feedback. We also thank McGill University’s Department of Bioengineering for the financial and scientific support that enabled the project. For the purpose of open access, the authors have applied a CC-BY license to this work and an MIT license to the associated software.

## AUTHOR CONTRIBUTIONS

HS: Conceptualization (supporting), Data curation (equal), Formal analysis (lead), Investigation (lead), Methodology (lead), Project administration, Software (lead), Validation (lead), Visualization (lead), Writing — original draft (lead), Writing — review & editing (lead).

DV: Data curation (equal), Formal analysis (supporting), Investigation (supporting), Methodology (supporting), Visualization (supporting), Writing — original draft (supporting).

VN: Formal analysis (supporting), Investigation (supporting), Software (supporting), Validation (supporting), Visualization (supporting), Writing — original draft (supporting).

ML: Formal analysis (supporting), Visualization (supporting), Writing — original draft (supporting).

JM: Data curation (supporting), Formal analysis (supporting), Visualization (supporting), Writing — original draft (supporting).

CK: Conceptualization (supporting), Resources, Supervision, Writing — review & editing (supporting).

SS: Conceptualization (supporting), Resources, Supervision, Writing — review & editing (supporting).

ME: Conceptualization (lead), Funding acquisition, Resources, Supervision, Writing — review & editing (supporting).

