## Supplemental Information for "PolyFold: Homology Filtering Approach Alters Apparent Structure Prediction Model Performances"

### Supplementary Information

#### A: Head-to-Head General Performance with a More Constrained Homology Cutoff of 0.4

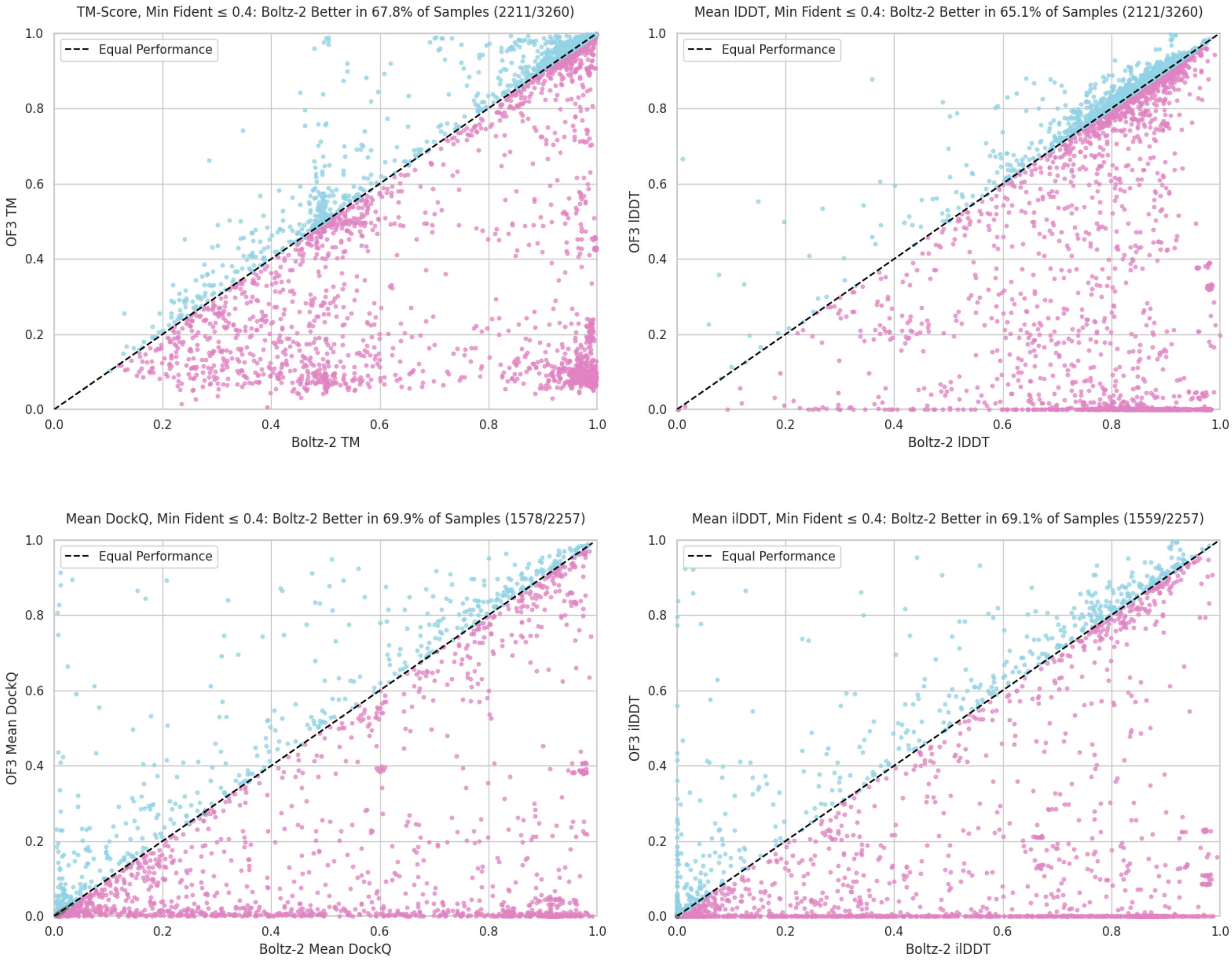


*Figure 1. General comparison of Boltz-2 vs. OpenFold 3 on (top left) TM-score, (top right) mean lDDT, (bottom left) mean DockQ, and (bottom right) mean ilDDT using a length-weighted average $F_{ident}$ cutoff over all chains in a structure as opposed to thresholding based solely on the chain with the lowest $F_{ident}$ in a structure. The rate of data points favouring Boltz-2 provided in each panel title is the count of data points below the dashed diagonal, representing the line of equal performance. Inclusion on the plot requires the structure's overall $F_{ident}$ ≤ 0.4 compared to the union of Boltz-2 and OpenFold 3 training sets.*

Minimum $F_{ident}$ must be less than or equal to 0.4 to be included in these plots.

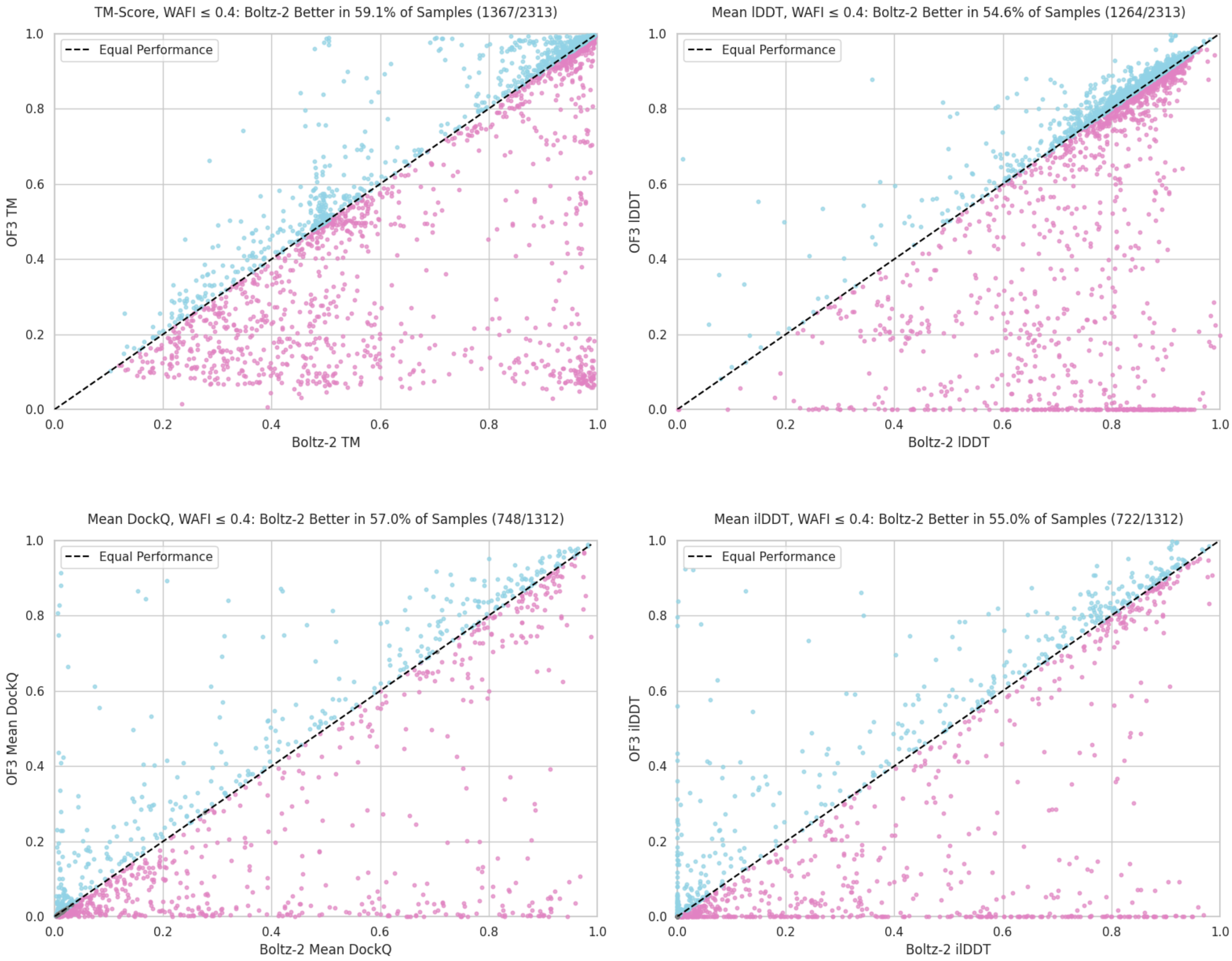


*Figure 2. General comparison of Boltz-2 vs. OpenFold 3 on (top left) TM-score, (top right) mean lDDT, (bottom left) mean DockQ, and (bottom right) mean ilDDT using a length-weighted average $F_{ident}$ cutoff over all chains in a structure as opposed to thresholding based solely on the chain with the lowest $F_{ident}$ in a structure. The rate of data points favouring Boltz-2 provided in each panel title is the count of data points below the dashed diagonal, representing the line of equal performance. Inclusion on the plot requires the structure's overall WAFI ≤ 0.4 compared to the union of Boltz-2 and OpenFold 3 training sets.*

WAFI must be less than or equal to 0.4 to be included in these plots.

#### B: Evaluation Set Data Partitioning

Table 1: Data Partitions and Metadata Extracted for Each Example

| **Column Use** | **Column Name** | **Meaning of Column** |
|---|---|---|
| PDB ID | pdb_id | Reference PDB identifier for the structure |
| Deposit Date | deposit_date | Date the structure was deposited to the PDB |
| Initial Release Date | initial_release_date | Date the structure was first released publicly |
| Experimental Method | experimental_method | Experimental technique used to determine the ground truth structure (e.g. cryo-EM, X-Ray, NMR) |
| Structure Resolution | resolution_combined | Structural resolution of the ground truth structure |
| File Size | file_size | Size of the raw CIF file on disk |
| Sequence Length | seq_length | Total length of all polymer chains (amino acids + nucleic acids) derived from FASTA and accounting for multimers |
| In Evaluation Set | in_eval_set | Binary indicator (1 = present in evaluation dataset, 0 = absent) |
| Polymer Entity Count | polymer_entity_count | Number of unique polymer entities in the CIF file (*not* total chains) |
| Nonpolymer Entity Count | nonpolymer_entity_count | Number of unique non-polymer entities in the CIF file |
| Assembly Count | assembly_count | Number of biological assemblies defined in the structure |
| Contains Small Molecule | contains_small_molecule_ligand | Presence of biologically relevant small molecules excluding solvents, glycans, and crystallization agents (see exclusion Table in Appendix B reproduced from the AlphaFold 3 paper) |
| Contains Ion | contains_included_ion | Presence of curated biologically relevant ions (e.g., $Mg^{2+}$, $Zn^{2+}$ from the inclusion Table in Appendix B reproduced from the AlphaFold 3 paper) |
| Modified Residues | contains_modified_residue_json | Presence of modified residues identified via custom parsing pipeline and stored as a JSON file which the workflow automation tool uses as an input when generating the prediction file. _pdbx_struct_mod_residue was parsed from the CIF file where available falling back to parsing _entity_poly.pdbx_seq_one_letter_code and ensuring to exclude glycans and canonical nucleotides |
| Enzyme Indicator | contains_enzyme | Binary indicator for enzyme presence based on EC annotation mapping |
| EC Codes | ec_code_by_chain | Enzyme Commission (EC) annotations mapped per chain |

| | | |
|---|---|---|
| Enzyme Chain Count | n_enzyme_chains | Number of chains annotated as enzymes |
| UniProt Accession | uniprot_acc | UniProt accession identifiers associated with the structure |
| Raw Localization | uniprot_loc_raw | Raw UniProt subcellular localization annotations |
| Localization | localization | Standardized localization category (membrane, cytosolic, secreted, unknown) |
| Protein Chains | protein_chain_count | Number of protein chains in the structure |
| Nucleic Acid Chains | nucleic_acid_chain_count | Number of nucleic acid chains |
| Has Nucleic Acid | has_nucleic_acid | Binary indicator for presence of nucleic acids |
| Structure Type | mer_type | Classification (monomer, dimer, multimer, protein-NA complex, etc.) |
| Peptide Chain Count | peptide_chain_count | Number of short peptide chains (<50 amino acids) |
| Minimum Peptide Length | peptide_min_chain_len | Length of shortest peptide chain |
| Has Peptide | has_peptide_chain | Binary indicator for presence of peptide chains |
| Is Peptide Structure | is_peptide | Binary indicator if structure is primarily peptide-based |
| Unique Protein Sequences | n_unique_protein_seqs | Number of distinct protein sequences across chains |
| Unique Nucleic Acid Sequences | n_unique_nucleic_acid_seqs | Number of distinct nucleic acid sequences across chains |
| Stoichiometry | protein_stoich | String representation of stoichiometry (e.g., A2B2) |
| Oligomer Type | oligomer_subtype | Classification as homomer or heteromer |
| Oligomer Reason | oligomer_reason | Explanation when classification is ambiguous |
| Antibody-like | is_antibody_like | Indicates antibody-like structures based on annotations and heuristics. Binary classification (1 = antibody-like, 0 = not antibody-like) |
| TCR Complex | is_tcr_complex | Indicates T-cell receptor complexes |
| MHC Complex | is_mhc_complex | Indicates MHC-containing complexes |
| Immune Class | immune_class | Aggregated immune-related classification |
| Fractional Identity | fident_n | Sequence identity for chain *n* from MMseqs2 alignment. Entries are organized by best hits, with the lowest *n* being the lowest $F_{ident}$ score |

| | | |
|---|---|---|
| Best Hit | best_hit_n | PDB ID of the best matching sequence from database |
| Hit Entity | hit_entity_n | To which chain of the PDB ID entity is the matching hit. This numerical annotation is generated via MMseqs2 |
| Chain Length | chain_n_length | Entity Length times the Multiplicity of Entity (see below). |
| Chain ID | chain_id_n | The chain ID for the specific chain in a given input file. |
| Entity Length | entity_n_length | Raw sequence length of the entity corresponding just to that one chain. |
| Multiplicity of Entity | multiplicity_n | This identifies how many times a particular entity appears in a FASTA. A homodimer would have an entity with multiplicity 2. |
| Weighted Average Fractional Identity | weighted_average_fident | Length-weighted average sequence identity across chains |
| WAFI Chain Count | waf_ready_chain_count | Number of chains included in WAFI calculation |
| Ligand RMSD | ligand_n_rmsd | RMSD of ligand *n* computed via OST |
| Ligand lDDT | ligand_n_lddt | lDDT score for ligand *n* |
| UniRef Neff | chain_X_uniref_neff | Effective sequence depth from UniRef for chain X |
| UniClust Neff | chain_X_uniclust_neff | Effective sequence depth from UniClust for chain X |
| Avg UniRef Neff | avg_unpaired_uniref_neff | Average UniRef $N_{eff}$ across chains |
| Avg UniClust Neff | avg_unpaired_uniclust_neff | Average UniClust $N_{eff}$ across chains |
| Avg Neff | avg_unpaired_neff | Combined average effective sequence depth |
| Superfamily ID | interpro_homologous_superfamily_id | InterPro homologous superfamily identifiers |
| Superfamily Name | interpro_homologous_superfamily_name | Names of homologous superfamilies |
| Family ID | interpro_family_id | Broadest InterPro family-level identifier |
| Family Name | interpro_family_name | Corresponding family name |
| Subfamily ID | interpro_subfamily_id | Most specific family-level identifier |
| Subfamily Name | interpro_subfamily_name | Corresponding subfamily name |
| Family Path IDs | interpro_family_path_ids | Full hierarchy of family IDs (broad → specific) |
| Family Path Names | interpro_family_path_names | Full hierarchy of family names |
| Domain IDs | interpro_domain_id | All matched InterPro domain identifiers |
| Domain Names | interpro_domain_name | Names of matched domains |

Beyond these data partitions, we also included OpenStructure metrics for both models in the final CSV.

#### C: Exclusion List of CCD Codes, and Accepted Ions

Table 2: Included Ions CCD code list

118, 119, 1AL, 1CU, 2FK, 2HP, 2OF, 3CO, 3MT, 3NI, 3OF, 4MO, 4PU, 4TI, 543, 6MO, AG, AL, ALF, AM, ATH, AU, AU3, AUC, BA, BEF, BF4, BO4, BR, BS3, BSY, CA, CAC, CD, CD1, CD3, CD5, CE, CF, CHT, CO, CO5, CON, CR, CS, CSB, CU, CU1, CU2, CU3, CUA, CUZ, CYN, DME, DMI, DSC, DTI, DY, E4N, EDR, EMC, ER3, EU, EU3, F, FE, FE2, FPO, GA, GD3, GEP, HAI, HG, HGC, HO3, IN, IR, IR3, IRI, IUM, K, KO4, LA, LCO, LCP, LI, LU, MAC, MG, MH2, MH3, MMC, MN, MN3, MN5, MN6, MO, MO1, MO2, MO3, MO4, MO5, MO6, MOO, MOS, MOW, MW1, MW2, MW3, NA2, NA5, NA6, NAO, NAW, NET, NI, NI1, NI2, NI3, NO2, NRU, O4M, OAA, OC1, OC2, OC3, OC4, OC5, OC6, OC7, OC8, OCL, OCM, OCN, OCO, OF1, OF2, OpenFold 3, OH, OS, OS4, OXL, PB, PBM, PD, PER, PI, PO3, PR, PT, PT4, PTN, RB, RH3, RHD, RU, SB, SE4, SEK, SM, SMO, SO3, T1A, TB, TBA, TCN, TEA, TH, THE, TL, TMA, TRA, V, VN3, VO4, W, WO5, Y1, YB, YBoltz-2, YH, YT3, ZCM, ZN, ZN2, ZN3, ZNO, ZO3, ZR

Table 3: Ligand Exclusion List of CCD Codes

144, 15P, 1PE, 2F2, 2JC, 3HR, 3SY, 7N5, 7PE, 9JE, AAE, ABA, ACE, ACN, ACT, ACY, AZI, BAM, BCN, BCT, BDN, BEN, BME, BO3, BTB, BTC, BU1, C8E, CAD, CAQ, CBM, CCN, CIT, CL, CLR, CM, CMO, CO3, CPT, CXS, D10, DEP, DIO, DMS, DN, DOD, DOX, EDO, EEE, EGL, EOH, EOX, EPE, ETF, FCY, FJO, FLC, FMT, FW5, GOL, GSH, GTT, GYF, HED, IHP, IHS, IMD, IOD, IPA, IPH, LDA, MB3, MEG, MES, MLA, MLI, MOH, MPD, MRD, MSE, MYR, N, NA, NH2, NH4, NHE, NO3, O4B, OHE, OLA, OLC, OMB, OME, OXA, P6G, PE3, PE4, PEG, PEO, PEP, PG0, PG4, PGE, PGR, PLM, PO4, POL, POP, PVO, SAR, SCN, SEO, SEP, SIN, SO4, SPD, SPM, SR, STE, STO, STU, TAR, TBU, TME, TPO, TRS, UNK, UNL, UNX, UPL, URE

Table 4: Crystallization Aids CCD Codes to Exclude

SO4, GOL, EDO, PO4, ACT, PEG, DMS, TRS, PGE, PG4, FMT, EPE, MPD, MES, CD, IOD

#### D: HDBSCAN Clustering with Effect Size & Multivariate Regression Analysis for Clusters

HDBSCAN clustering was performed on both a 10D vector and a 20D with min_cluster_size=20. The 10D vector simply did not include: Boltz-2_ligand_avg_rmsd, Boltz-2_ligand_avg_lddt, Boltz-2_model_clashes, Boltz-2_bad_angles, and Boltz-2_bad_bonds (same for OpenFold 3_*). It only included the other metrics mentioned in the methods (RMSD, DockQ, TM, lDDT, and iLDDT). We performed dimensionality reduction of our clustering to determine if distinct clusters were formed. Additionally, we plotted the metrics corresponding with each cluster to visually analyze the separation of clusters.

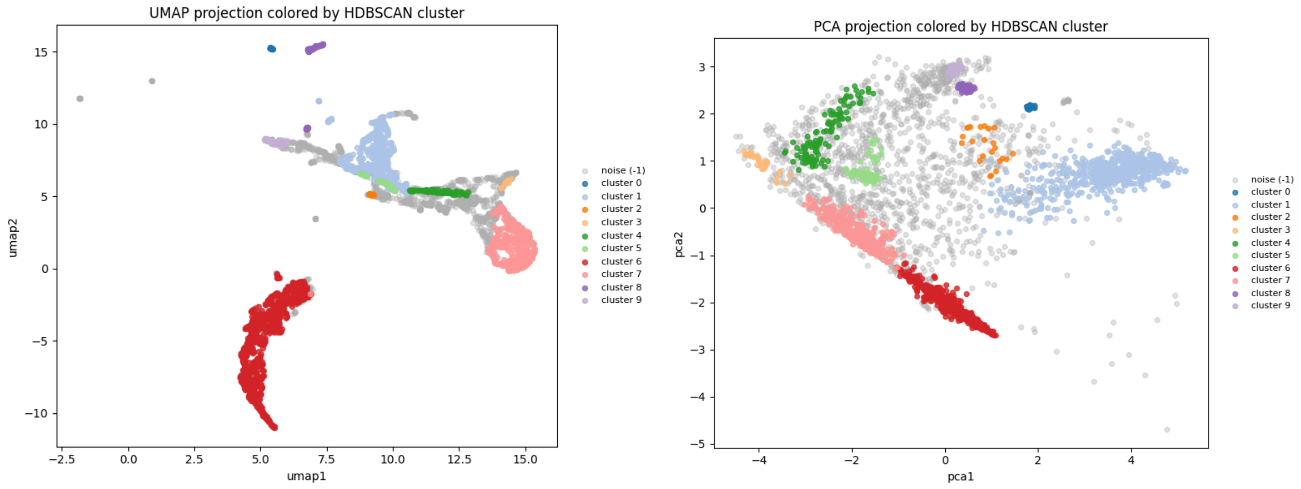


*Figure 3: Dimensionality reduction using UMAP (left) and PCA (right) of HDBSCAN clustering performed on our 10-dimensional vectors.*

Dimensionality reduction was performed using PCA and UMAP to verify that there are distinct clusters in the data. We tested both methods because we were not certain the relationships would be linear (where PCA would work), so we wanted to ensure there was clear clustering with both dimensionality reduction methods (UMAP would have captured non-linear dimensionality reduction as well).

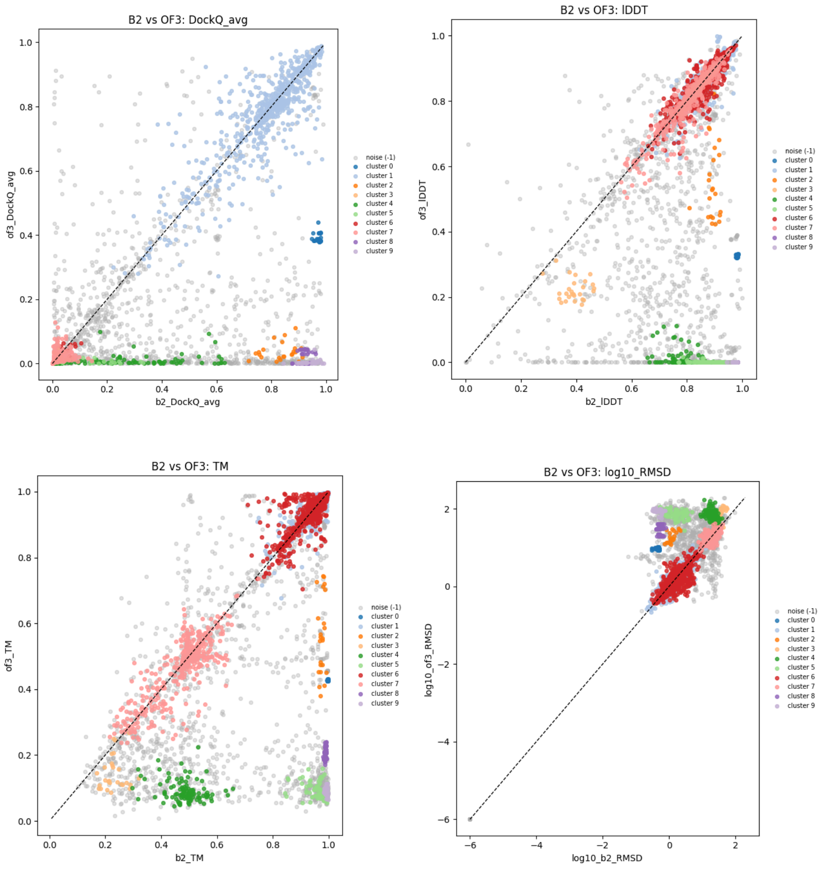


*Figure 4: 10D clusters assigned to head-to-head metrics (DockQ, lDDT, TM, RMSD).*

We performed the same dimensionality reduction and plotting of our clusters on our 20D vector and noticed that this generated fewer clusters. Largely, the clusters were similar across the 10D and 20D clustering, however, there were a couple of clusters that existed for 10D clustering which did not for 20D clustering.

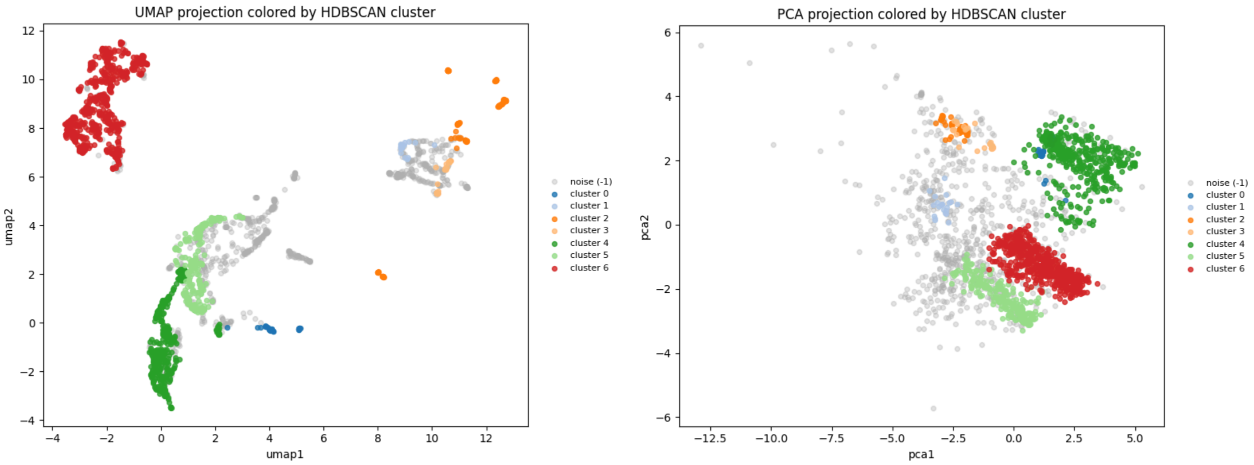


*Figure 5: Dimensionality reduction using UMAP (left) and PCA (right) of HDBSCAN clustering performed on 20-dimensional vectors.*

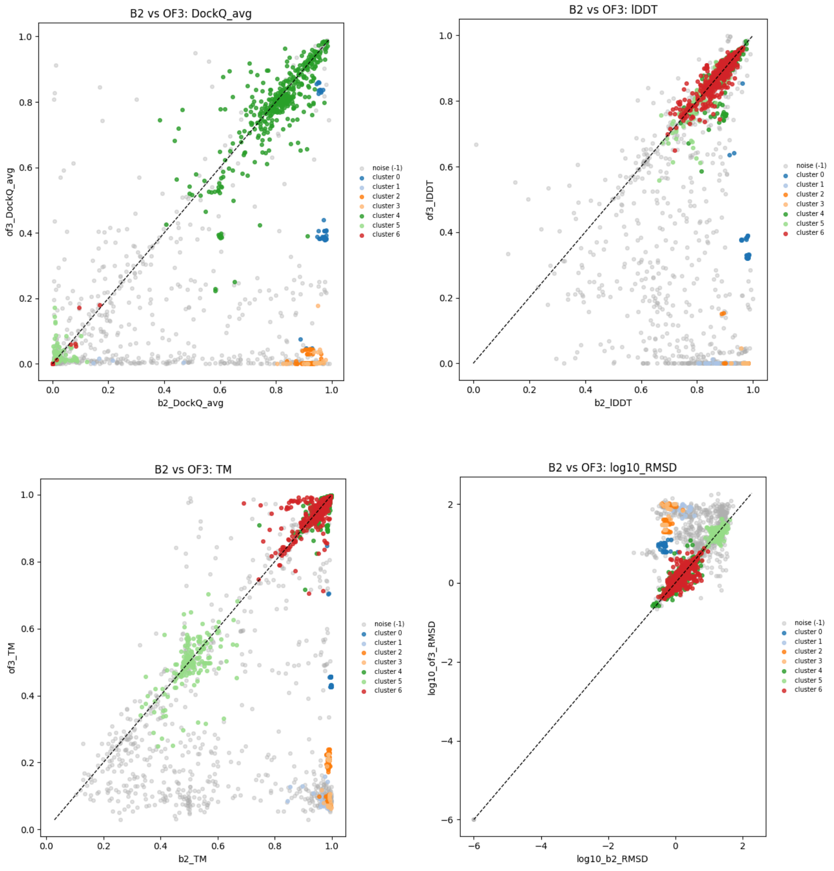


*Figure 6: 20D clusters assigned to head-to-head metrics (DockQ, lDDT, TM, RMSD).*

Interestingly, across both dimensions we notice clusters having extremely low RMSD for both Boltz-2 and OpenFold 3. When investigating these outliers closer, we noticed that generally they have very high WAFI which suggests that they are largely residual homology leakage. All contain traces of residual homology leakage to structures released before 2021-09-30, meaning both models had access to these structures, which explains their abnormally strong performance (PDB ID: 4N14 has residual homology leakage from both models). There are also some structures which have a WAFI of 0; these are all Cryo-EM structures of intermolecular π-electron interactions within π-conjugated peptidic supramolecular polymers (PDB ID: 9O7J and 9O7L).

To evaluate the differences in clusters to the rest of the data, we performed effect size analysis to compare which inputs made a cluster different from all the other data points, and a multivariate regression to determine what the most significant driver of performance within the cluster was (see Methods in the main work).

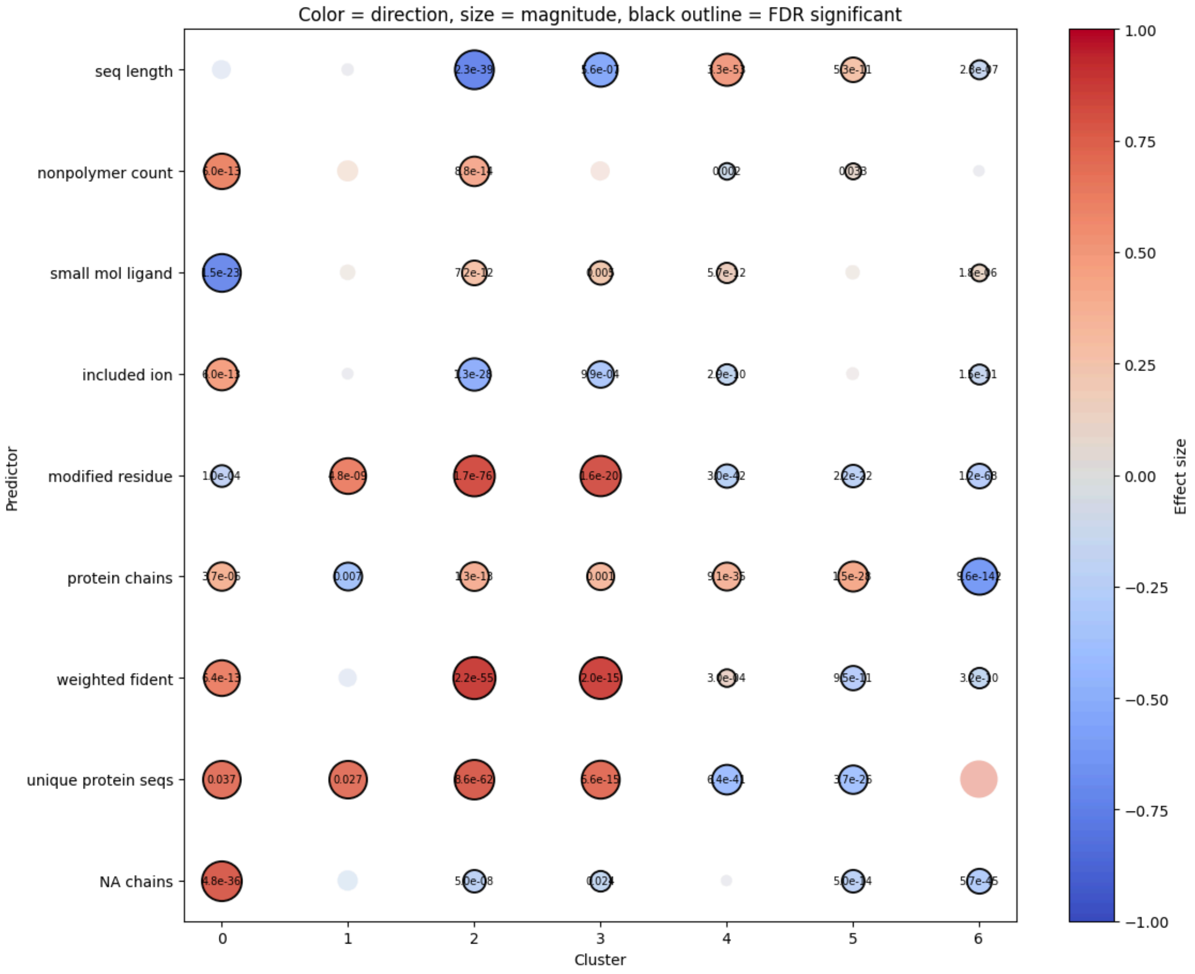


*Figure 7: Cluster-level predictor enrichment and depletion for 20D clustering . Bubble plot summarizing effect sizes for each predictor across clusters (one-vs-rest comparisons). Color indicates direction of effect (red: enriched/higher in cluster; blue: depleted/lower), while bubble size reflects the magnitude of the effect. Black outlines denote predictors that are statistically significant after Benjamini–Hochberg false discovery rate (FDR) correction (q < 0.05). Effect sizes correspond to rank-biserial correlation for continuous variables and proportion differences for binary variables.*

We see that there are multiple inputs which make a cluster distinct from the rest of the data, and these are all aspects we looked at manually when inspecting the clusters. The effect size was helpful in confirming that the effect we were seeing was indeed statistically significant. We additionally performed the same analysis across the 10D clustering.

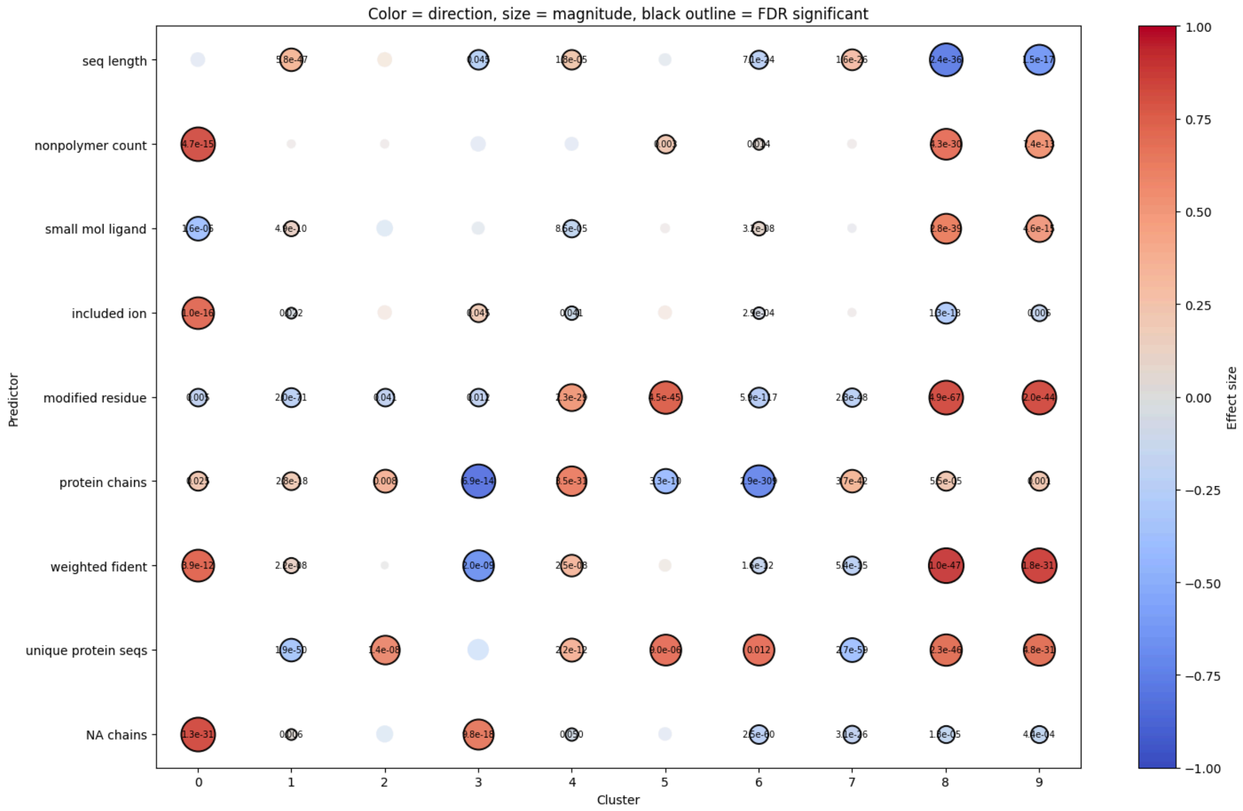


*Figure 8: Cluster-level predictor enrichment and depletion for 10D clustering. Bubble plot summarizing effect sizes for each predictor across clusters (one-vs-rest comparisons). Color indicates direction of effect (red: enriched/higher in cluster; blue: depleted/lower), while bubble size reflects the magnitude of the effect. Black outlines denote predictors that are statistically significant after Benjamini–Hochberg false discovery rate (FDR) correction (q < 0.05). Effect sizes correspond to rank-biserial correlation for continuous variables and proportion differences for binary variables.*

#### Multivariate regression

Multivariate regression was performed as described across the Methods section in the main work. This resulted in coefficients across all 5 metrics (RMSD, TM, lDDT, iLDDT, and DockQ) across every cluster, so for brevity, we only show lDDT and TM for cluster 4 as an example here.

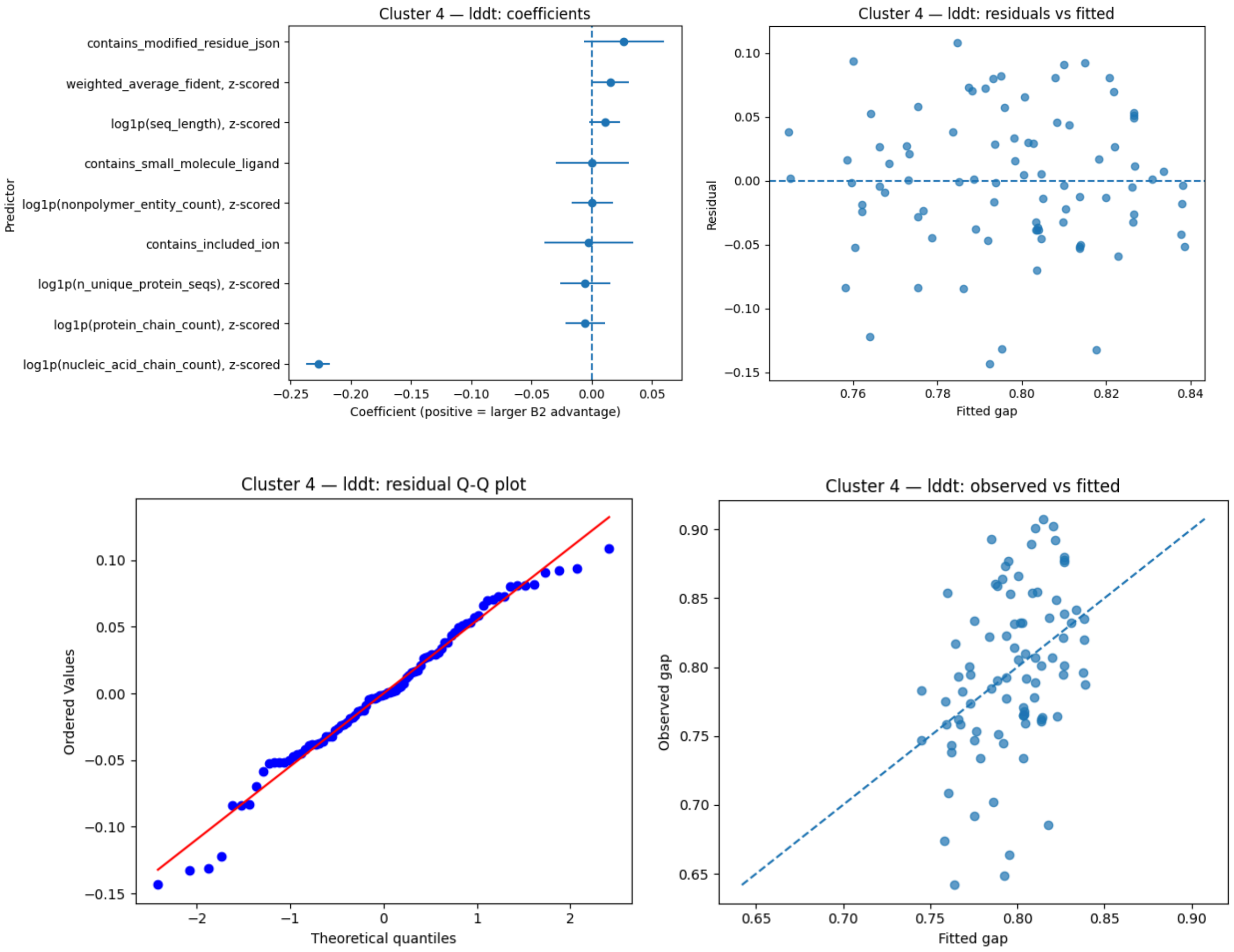


*Figure 9: Cluster 4 specific multivariate regression analysis of model performance differences in lDDT. Linear regression coefficients for Cluster 4 are shown for the selected performance metric, with error bars representing 95% confidence intervals estimated using HC3 robust standard errors. Positive coefficients indicate predictors associated with improved performance of Boltz-2 relative to OpenFold 3, whereas negative coefficients indicate the opposite. Predictors were log-transformed and standardized where appropriate prior to modeling. Model diagnostics are shown on the right: residuals vs. fitted values demonstrate no strong systematic bias, the Q-Q plot indicates approximate normality of residuals, and the observed vs. fitted comparison illustrates overall model fit.*

We generate the same coefficients information, residuals vs. fitted, residuals Q-Q plots, and observed vs. fitted plots across all clusters for the 5 metrics outlined in Methods. See below for the rest of the data for cluster 4.

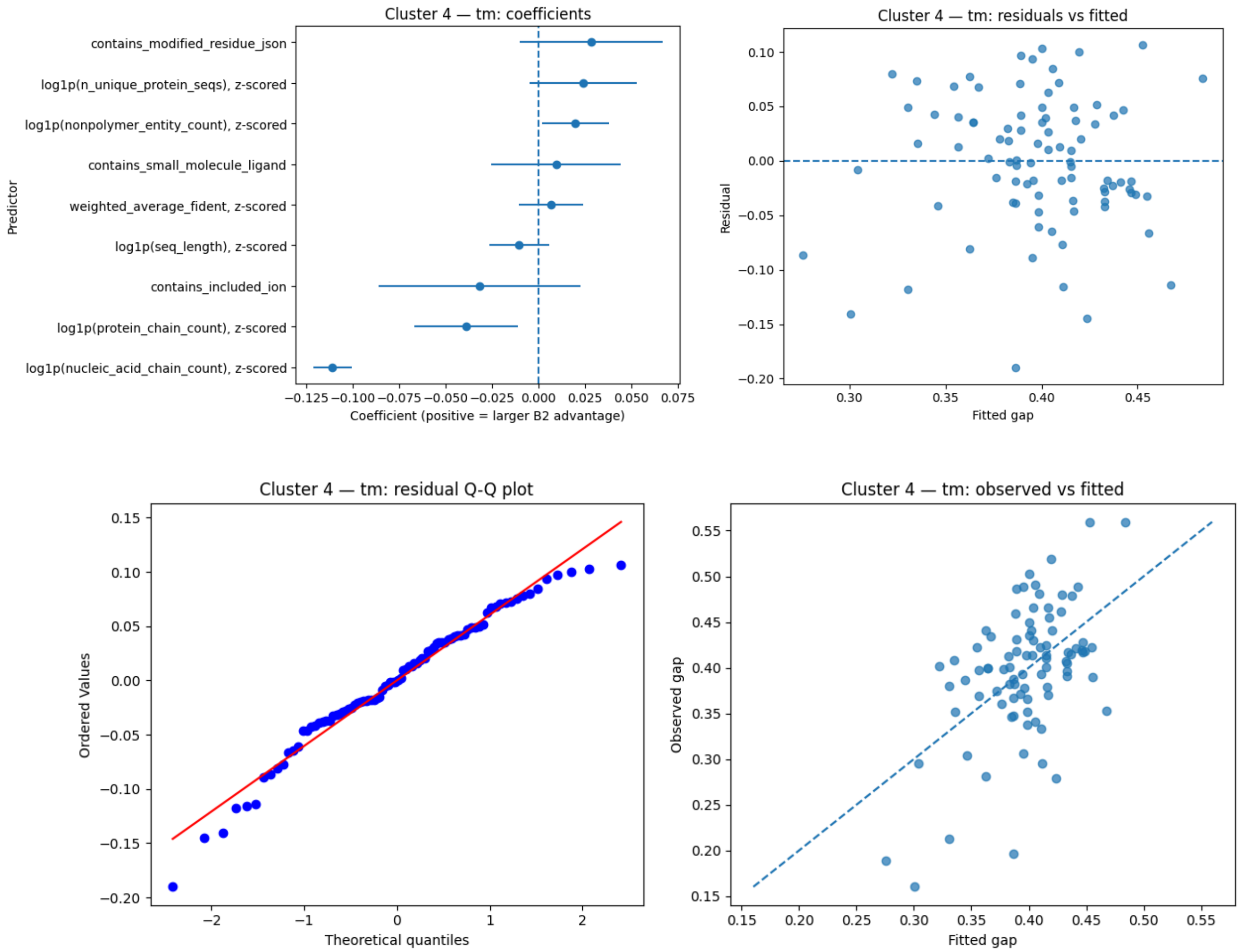


*Figure 10: Cluster 4 specific multivariate regression analysis of model performance differences in TM. Linear regression coefficients for Cluster 4 are shown for the selected performance metric, with error bars representing 95% confidence intervals estimated using HC3 robust standard errors. Positive coefficients indicate predictors associated with improved performance of Boltz-2 relative to OpenFold 3, whereas negative coefficients indicate the opposite. Predictors were log-transformed and standardized where appropriate prior to modeling. Model diagnostics are shown on the right: residuals vs. fitted values demonstrate no strong systematic bias, the Q-Q plot indicates approximate normality of residuals, and the observed vs. fitted comparison illustrates overall model fit.*

Different metrics have different coefficients which are more significant than others. These are helpful to rapidly observe if there are any inputs (as defined in the clustering subsections in Methods in the main work) which consistently affect the performance of one model over another in a given cluster.
